# Highly reproducible 16S sequencing facilitates measurement of host genetic influences on the stickleback gut microbiome

**DOI:** 10.1101/497792

**Authors:** Clayton M. Small, Mark Currey, Emily A. Beck, Susan Bassham, William A. Cresko

## Abstract

Multicellular organisms interact with resident microbes in important ways, and a better understanding of host-microbe interactions is aided by tools such as high-throughput 16S sequencing. However, rigorous evaluation of the veracity of these tools in a different context from which they were developed has often lagged behind. Our goal was to perform one such critical test by examining how variation in tissue preparation and DNA isolation could affect inferences about gut microbiome variation between two genetically divergent lines of threespine stickleback fish maintained in the same lab environment. Using careful experimental design and intensive sampling of individuals, we addressed technical and biological sources of variation in 16S-based estimates of microbial diversity. After employing a two-tiered bead beating approach consisting of tissue homogenization followed by microbial lysis in subsamples, we found an extremely minor effect of DNA isolation protocol relative to among-host microbial diversity differences. Individual abundance estimates for rare OTUs, however, showed much lower reproducibility. We found that the stickleback gut microbiome was highly variable, even among siblings housed together, but that an effect of host genotype (stickleback lineage) was detectable for some microbial taxa. Our findings demonstrate the importance of appropriately quantifying biological and technical variance components when attempting to understand major influences on high-throughput microbiome data.

## INTRODUCTION

From early development through senescence, animal and plant hosts interact with their resident microbiota through complex host-microbe relationships, resulting in a diversity of both positive and negative outcomes for host health and fitness. Beneficial microbes prime ontogenesis of immunity (Iatsenko *et al.* 2014; Sudo *et al.* 1997), regulate the host inflammatory response (Olszak *et al.* 2012), and aid in various digestive and metabolic processes (Hooper *et al.* 2001; Russell & Rychlik 2001). Pathogenic microbes release toxins (Simon *et al.* 2014), disrupt microbial community structure (Gillis *et al.* 2018), and may contribute to human diseases such as diabetes, obesity, and inflammatory bowel disease (IBD), for example (Wu *et al.* 2015). The complexity of host-microbe interactions, and significant consequences for host wellness when the relationships are perturbed, argue that many host-microbe relationships are, at least in part, a result of co-evolution. For example, the intimate symbiotic relationship between Hawaiian bobtail squids and the luminescent bacterium *Vibrio fischeri* (McFall-Ngai *et al.* 2012; McFall-Ngai & Ruby 1991), presents a strong case for evolutionary novelty arising from co-evolution. Indeed, the recognized importance of host-microbe interactions has led to a recent spike in interdisciplinary research efforts, complete with accelerated tool development both molecular and computational in nature. This rapid progress, however, has in some cases meant a lag in the thorough evaluation of the veracity and efficacy of these tools.

Understanding host-microbe relationships from ecological, evolutionary, and disease perspectives hinges on quantification of microbial diversity in samples from various host body sites. Although quantification is increasingly being achieved using shotgun metagenomic approaches (Qin *et al.* 2012; Sharpton 2014), marker-based techniques such as high-throughput 16S rRNA amplicon sequencing are still the most cost-effective, straightforward, and commonly applied methods for microbial community profiling. In the last decade, high-throughput 16S sequencing has expanded from applications in environmental, human, and laboratory model host contexts, to uses for a variety of truly diverse host plant and animal sample types (Hyde *et al.* 2016; Nuccio *et al.* 2016; Torres *et al.* 2017).

As researchers extend their work beyond routinely characterized environments such as soil and human fecal samples and into new, diverse study systems, the adoption and extension of previously optimized techniques should occur cautiously and intentionally. Methodologies for 16S amplicon sequencing should ideally be evaluated at multiple stages (i.e. sample collection and handling through analysis), compared with multiple alternative options, and evaluated with respect to the discriminatory power and precision of diversity analyses based on them. The Microbiome Quality Control Project (Sinha *et al.* 2017), for example, has addressed some of these issues for human stool and artificial microbial communities, including an effective quantification of laboratory-to-laboratory variation. Other diverse endeavors have evaluated effects of DNA isolation attributes on sequencing-based community inference in corals (Weber *et al.* 2017), fleas (Lawrence *et al.* 2015), human saliva (Raju *et al.* 2018), and marine biofilms (Corcoll *et al.* 2017), for example, but the foci of these studies did not include quantifying reproducibility and its uncertainty using large samples of among-individual variation.

Research aims may require the direct sampling of whole host organs in animal models, such as the gastrointestinal tract, to obtain an unfiltered view of internal host-microbe relationships. However, with these shifts in sampling strategy comes a slew of considerations and obstacles, in part because commercially available kits are designed and optimized for a narrow range of sample types such as soil or human stool. Potential problems with sample processing and library preparation based on these unrefined protocols may include poor DNA integrity, inadequate DNA quantity, host and reagent contamination, and low repeatability, all of which may vary depending on the sample type.

We compared the gut microbiomes of two divergent populations of threespine stickleback (*Gasterosteus aculeatus*) that have been maintained in the same lab environment to evaluate whether our biological conclusions could be affected by the use of three popular DNA isolation protocols. Threespine stickleback fish have repeatedly colonized a diversity of freshwater habitats from ancestral marine populations, resulting in exceptional degrees of within- and among-population genetic and phenotypic variation for countless traits (Bell & Foster 1994; Colosimo *et al.* 2004; Cresko *et al.* 2004; Cresko *et al.* 2007; Glazer *et al.* 2015; Hohenlohe *et al.* 2010). Genetic diversity among stickleback populations is in many ways similar to variation in humans, making stickleback an excellent model for understanding the role of host genetic variation in determining phenotypes germane to host-microbe interactions, including microbial community structure itself (Milligan-Myhre *et al.* 2016; Small *et al.* 2017; Smith *et al.* 2015).

In our initial attempts to isolate DNA from adult stickleback guts for 16S sequencing we found commonly used DNA isolation protocols, including one specifically designed for microbial samples, untenable owing to low quality and high variance of DNA yield, and fragmentation both within and among protocols. This outcome prompted us to optimize these DNA isolation protocols for adult stickleback guts.

We employed careful experimental design and thorough sampling of laboratory-raised hosts to address both technical and biological sources of variation in 16S-based estimates of microbial diversity. The relative contributions of individual host and DNA isolation protocol to variation in 16S-based diversity estimates have not been satisfactorily measured in previous studies, due to insufficient biological (individual-level) replication, inadequate parameter estimation, or both. To address this, the technical objectives of our study included a careful comparison of operational taxonomic unit (OTU) relative abundance and diversity (both alpha and beta) across libraries generated from three DNA isolation protocols, followed by formal quantification of reproducibility and its uncertainty. The biological objective of our study was to test for differences in relative OTU abundance and diversity arising from genetic differences between two stickleback laboratory lines, one descended from a freshwater lake population and the other from an oceanic population, but both raised and housed in a common environment. The factorial nature of our study design also permitted assessment of statistical interactions between DNA isolation protocol and host genotype, that is, whether any dependency of biological inferences on DNA isolation protocol choice might exist. We also performed a separate experiment in which we measured the precision of each DNA isolation protocol using replicate samples from the same individual.

In general we found the stickleback gut microbiome to be highly variable even among individuals of the same sibship, and that variation due to the two host genetic backgrounds (population of origin) was detectable but smaller than individual-level variation. Unoptimized tissue processing had a major effect on the yield and integrity of DNA isolated using different protocols. However, after employing a two-step bead beating approach consisting of initial tissue homogenization followed by microbial lysis in homogenate subsamples, we found an extremely minor effect of DNA isolation protocol on the ability to understand microbial diversity using 16S data. This is an important finding for those researchers faced with the decision of having to choose among available protocols, so long as they take necessary measures to reduce bias during initial tissue processing steps. Our protocol optimization, study design, and insights both technical and biological should be useful to others who seek to quantify microbial community structure in fish guts and other tissue types using high throughput 16S sequencing.

## MATERIALS AND METHODS

### Rearing of adult stickleback, evaluation of DNA quality, and experimental design

#### Threespine stickleback husbandry and collection of gut samples

We collected guts from male adult threespine stickleback (*Gasterosteus aculeatus*) derived from wild-caught Alaskan populations, which have been maintained in the laboratory for at least 10 generations. All individuals were raised to an age of 12 to 16 months, using standard protocols described in previous publications (Cresko *et al.* 2004). Briefly, fish were raised from embryos fertilized in vitro, and larvae were fed twice daily with brine shrimp nauplii and Zeigler Larval AP100 Diet. An equal parts mixture of Golden Pearl 800-1000 Micron Juvenile Diet, Otohime C1, Zeigler Zebrafish Diet, and Hikari Tropical Micro Pellets was fed twice daily to fish as juveniles and adults. Fish were housed in a large recirculating system with a 10% daily water change, in 20-L tanks at a density of 20-30 fish per tank. Tanks were randomly positioned on a single shelving rack roughly equidistant from the incoming water source. We maintained fish in an approximately 1:1 sex ratio, with a photoperiod of 8 hours light and 16 hours dark (including 30 min. dawn and dusk).

To reduce among-individual variation owing to sex (Bolnick *et al.* 2014), we sampled males only, as confirmed by DNA isolation from caudal fin clips and a PCR-based sex genotyping procedure (see Small *et al.* 2017). Fish were also not fed for 24 hours prior to sampling to reduce the amount of food in the gut. Upon euthanasia by lethal dose of MS222, the entire gastrointestinal tract of each fish, including the esophagus to just anterior of the urogenital opening, was carefully removed, weighed, and quickly flash frozen in liquid nitrogen in a screw-top tube containing nuclease-free homogenization beads (see below).

#### Initial assessment of DNA quality from three unmodified DNA isolation protocols

We evaluated yield and quality of DNA isolated using a standard phenol-chloroform-isoamyl alcohol protocol (“PCP”) and two commercial kit protocols commonly used in microbiome studies: MO BIO’s PowerFecal^®^ Kit (“PFP”) and the Qiagen’s DNeasy Blood and Tissue Kit (“DEP”). We dissected whole guts from 52 adult stickleback in our fish facility, randomly assigning 20 each to PCP and PFP, and 12 to DEP. In this initial assessment of unmodified DNA isolation protocols, we used the entire gut to be consistent with previous studies of stickleback gut microbiota (Bolnick *et al.* 2014; Smith *et al.* 2015). In the case of PCP and DEP, each whole gut was dissected and flash frozen (see above) in a tube containing five nuclease-free 3.2-mm stainless steel beads (Next Advance, SSB32). In the case of PFP each gut was frozen in a tube containing ~1-mm garnet beads, which are standard issue for the kit. Next, we removed each tube containing a gut and beads from -80 C and followed manufacturer recommendations, with a few exceptions. We added 400 *μ*L of Qiagen Buffer ATL in the case of PCP and DEP, and 750 *μ*L of Bead Solution in the case of PFP. We then homogenized guts in a Thermo Savant FastPrep FP120 with three 40-second bouts of beating at intensity level 6.5. Using the entire homogenate for each sample, we followed instructions in the manuals of DEP and PFP kits. In the case of PCP, we followed the same post-homogenization lysis instructions as for DEP, then combined a 650-*μ*L aliquot of the lysate with 650 *μ*L of 25:24:1 equilibrated phenol-chloroform-isoamyl alcohol in a phase-lock gel tube, mixed by inversion, and centrifuged at 18,000 x*g* in a bench-top microcentrifuge for 5 minutes. We transferred the aqueous layer to a new phase-lock gel tube, added 500 *μ*L of 24:1 chloroform-isoamyl alcohol, mixed by inversion, centrifuged again, and transferred the aqueous layer to a 1.5-mL tube. We precipitated the DNA using 450 *μ*L of isopropanol and 5-minute centrifugation at 5,800 xg, washed the pellet once with 70% ethanol and twice with 95% ethanol, air-dried the pellets for 10-15 minutes, and resuspended the pellet in 100 *μ*L of Qiagen Buffer EB. We quantified DNA resulting from all three protocols using a Qubit 2.0 fluorometer (Invitrogen) and evaluated fragment length distributions using a Fragment Analyzer (Advanced Analytical).

#### Design of experiments

After optimizing these protocols (see below), we designed two separate experiments to evaluate effects of several variables on inferred microbial community structure. For these experiments we used two entirely different sets of fish sampled from our fish facility at different times, and obtained multiple subsamples per fish gut. For the first experiment each of 36 individual intestinal tracts were homogenized, then divided into three separate sub-samples to be analyzed using different DNA isolation protocols. This design effectively allowed us to measure among-protocol technical variation based on within-host comparisons (reproducibility). These 36 fish were from two different lab lines, specifically a freshwater (FW) line derived from the natural population “Boot Lake,” and an oceanic (OC) line derived from the natural population “Rabbit Slough.” We sampled three different full-sib families from each line, and six fish per family. Each family was housed in a different tank but in the same recirculating system. Our study design therefore enabled an assessment of the influence of host genetic background on the microbiota, a topic of great interest for the field of host-microbe interactions (Hildebrand *et al.* 2013; Kurilshikov *et al.* 2017; Rothschild *et al.* 2018). Fig 1A illustrates these components of the experimental design. Finally, using a sample of six FW males from a seventh full-sib family, we sampled two guts for each protocol type (six guts total), but we repeated six measurements (six subsamples) per homogenized gut to evaluate the within-fish repeatability (precision) of each DNA isolation protocol. Fig 1B reflects the precision assessment inherent in our study design.

**Figure 1.**
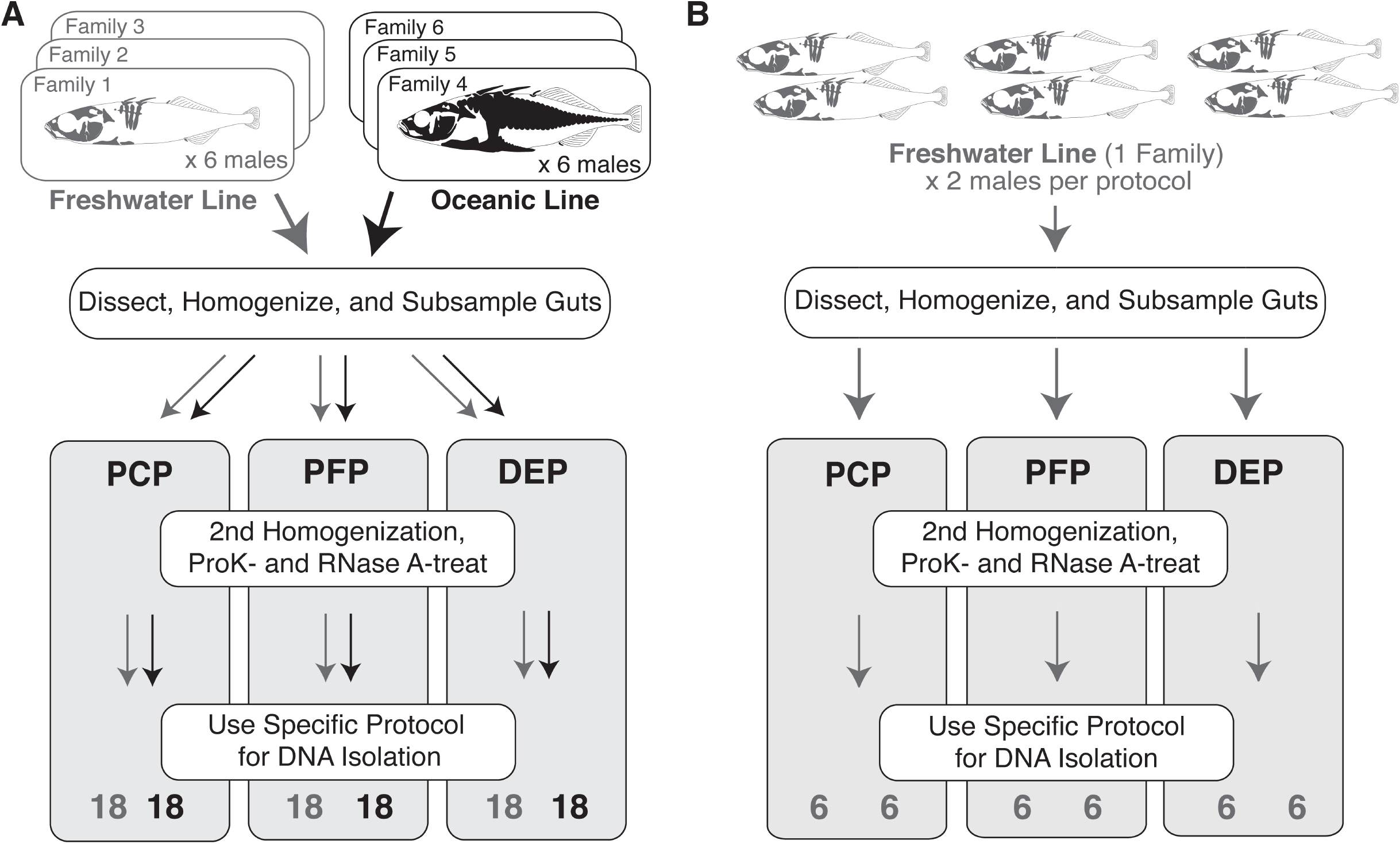
Experimental design to evaluate **A**. Technical (DNA isolation protocol) and biological (individual and population) variation in 16S sequencing-based diversity metrics, and **B**. Within-individual precision for these metrics. Shown are the stickleback lines (populations), families, and sample processing steps, and sample sizes used in the current study. In the first experiment (**A**), we assigned one of three homogenate subsamples from each fish gut to one of three DNA isolation protocols: Phenol-chloroform (PCP), PowerFecal (PFP), or DNeasy (DEP). In the second experiment (**B**), we assigned all six homogenate subsamples from a given fish gut to one of the DNA isolation protocols, with each protocol represented by two fish.

### Modified DNA isolation methods and Illumina 16S amplicon sequencing

#### Standardized pre-processing with 20 mg subsampling

In order to effectively compare the three DNA methods using our experimental design, we standardized tissue pre-processing and introduced uniform-mass tissue subsampling for all gut samples. We removed each gut (in a screw-cap tube with five nuclease-free 3.2-mm stainless steel beads) from -80 C, added 800 *μ*L of pre-warmed Qiagen Buffer ATL (with 0.5 *μ*M EDTA), and homogenized using the FastPrep FP120. We applied three bouts of 40-second beating at intensity level 6.5 to achieve a homogeneous mixture, then pipetted volumes from each sample to achieve 20-mg subsamples, as calculated from the original mass of each gut. For the reproducibility experiment (Fig 1A), three subsamples from each gut were taken, one for each of the 3 DNA isolation protocols described below. For the repeatability experiment (Fig 1B), six subsamples from each gut were taken, all for a single DNA isolation protocol. Subsamples were transferred to screw-top tubes containing 100 *μ*L of 0.15 mm zirconium oxide beads (NextAdvance, ZrOB015), for future mechanical lysis of microbes, flash frozen in liquid nitrogen, and stored at -80 C. These aliquots then received one of the 3 DNA isolation treatments below. We also performed two “negative control” DNA isolations for each isolation protocol, in which the protocol was carried out starting with no gut, and with or without the addition of proteinase K (see below). Because our objectives did not include comparisons of accuracy among DNA isolation protocols, as others (Salter *et al.* 2014; Yuan *et al.* 2012) have evaluated this, we did not incorporate controlled assemblages of microbes (“mock communities”) in our experimental design.

#### Phenol-chloroform-isoamyl alcohol protocol (PCP)

We removed homogenate subsamples from -80 C and added pre-warmed Qiagen Buffer ATL to bring the total ATL volume in the tube to 676 *μ*L. We then homogenized for two 40-second bouts in the FastPrep FP120 at level 6.5, briefly spun tubes, added 20 *μ*L of proteinase K (20 mg/mL), mixed by aspiration, and incubated at 56 C for 30 minutes. We added 4 *μ*L of RNase A (100 mg/mL), mixed by aspiration, incubated at 37 C for 30 minutes, and then transferred the entire volume of lysate to a new 1.5 mL tube, to which 500 *μ*L of 25:24:1 equilibrated phenol-chloroform-isoamyl alcohol was added. At this point we carried out the remainder of the phenol-chloroform protocol exactly as described above.

#### MoBio PowerFecal protocol (PFP)

We removed homogenate subsamples from -80 C and added pre-warmed PowerFecal Bead Solution to bring the total volume of solution (ATL + Bead Solution) in the tube to 750 *μ*L. We then added 60 *μ*L of PowerFecal C1 solution, incubated at 65 C for 10 minutes, and then homogenized for two 40-second bouts in the FastPrep FP120 at level 6.5. We briefly spun tubes, added 20 *μ*L of proteinase K (20 mg/mL), mixed by aspiration, and incubated at 56 C for 30 minutes. Then we added 4 *μ*L of RNase A (100 mg/mL), mixed by aspiration, incubated at 37 C for 30 minutes, and followed the instructions in the PowerFecal manual, starting with “step 7,” which is a centrifugation for 1 minute at 13000 x *g* to pellet and remove solids from the lysate. Finally, we eluted with 50 *μ*L Qiagen Buffer EB and quantified DNA concentration as above.

#### Qiagen DNeasy protocol (DEP)

We removed homogenate subsamples from -80 C and added pre-warmed Qiagen Buffer ATL to bring the total ATL volume in the tube to 776 *μ*L. We then homogenized for two 40-second bouts in the FastPrep FP120 at level 6.5, briefly spun tubes, added 20 *μ*L of proteinase K (20 mg/mL), mixed by aspiration, and incubated at 56 C for 30 minutes. We added 4 *μ*L of RNase A (100 mg/mL), mixed by aspiration, incubated at 37 C for 30 minutes. We combined the entire volume of lysate with 800 *μ*L of Qiagen Buffer AL and 800 *μ*L of 100% ethanol to a new 15 mL screw-cap tube to ensure adequate volume for the additional reagents. After briefly mixing by aspiration we transferred 600 *μ*L of the mixture to a DNeasy spin column, spun at 6,000 xg in a bench top microcentrifuge for 1 minute, discarded flow-through, and repeated four times, for the remainder of the mixture. Next we added 500 *μ*L of Qiagen solution AW1, spun at 6,000 xg for 1 minute, added 500 *μ*L of Qiagen solution AW2, spun at 18,000 x*g* for 3 minutes, added 500 *μ*L of 80% ethanol, and spun at 18,000 xg for 3 minutes. Finally, we eluted DNA with 100 *μ*L Buffer EB and quantified DNA concentration as above.

#### Construction of 16S rRNA gene amplicon libraries and Illumina sequencing

We submitted a 25 ng/*μ*L dilution from each gut DNA sample to the University of Oregon Genomics and Cell Characterization Core Facility (GC3F) for library amplification, cleanup and sequencing. All six negative control samples were not diluted, as the DNA in these samples was lower than the detection limit of our fluorometer. The GC3F generated 16S libraries from 200 ng of DNA template per sample using custom primers 515F (5’AAT GAT ACGGCGACCACCGAGAT CT ACACxxxxxxxxTATGGTAATTGTGTG CCAGCMGCCGCGGTAA3’) and 806R (5’CAAGCAGAAGACGGCAT ACGAGAT xxxxxxxxAGT CAGT CAGCCGGACT ACH VGGGTWTCTAAT3’), which are based on those described in Caporaso et al. (2011) and which amplify the “V4” 16S region but enable dual indexing (indexes represented by “x”s in the above sequences). A cocktail including 12.5 *μ*L NEBNext^®^ Q5^®^ Hot Start HiFi PCR Master Mix, 4.5 *μ*L of 2.79 *μ*M primer mix, and 8 *μ*L of DNA template, was used for each library PCR. The thermal profile was as follows: initial denaturation at 98 C for 30 seconds, followed by 22 cycles of 98 C for 10 seconds, 61 C for 20 seconds, and 72 C for 20 seconds, followed by final extension at 72 C for 2 minutes. Each library was cleaned twice using 20 *μ*L of Omega Mag-Bind^®^ RxnPure Plus beads and quantified by a Qubit fluorometer, at which point 9.235 ng of DNA from each library (less for negative controls) were pooled. The GC3F quantified the library pool using qPCR, combined it with a complex RNA-Seq library from an unrelated project, and sequenced 161-nt paired-end reads in two Illumina HiSeq 2500 lanes.

### Processing of Illumina 16S data and statistical inference

#### Sequence filtering and OTU picking

Processing of sequences and OTU picking was primarily achieved using accessory scripts from QIIME version 1.9.1 (Caporaso *et al.* 2010b) and to a lesser extent our own custom scripts. We overlapped ends of read pairs using QIIME’s join_paired_ends.py, and we demultiplexed the merged reads using QIIME’s extract_barcodes.py and split_libraries_fastq.py. We used default arguments, except that we allowed a maximum of two barcode errors when demultiplexing and invoked read truncation at 30 or more consecutive low quality base calls. This filtering process yielded 119.94 million total reads from the 144 gut libraries, and 4957 total reads from the six negative control libraries. We performed open reference OTU picking using QIIME’s pick_open_reference_otus.py with default settings (Caporaso *et al.* 2010a; Edgar 2010), which uses the Greengenes version 13.8 database as its reference (DeSantis *et al.* 2006). We then removed all OTUs of mitochondrial and chloroplast origin to exclude the influence of host- and food-derived DNA. The total number of filtered OTU-assigned reads from gut libraries was 87.109 million (mean = 604920.326; SEM = 26291.498). Negative control libraries produced exceedingly small numbers of OTU-assigned reads (PCP: 418; PCP_proK: 348; PFP: 1201; PFP_proK: 1115; DEP: 644; DEP_proK: 367). Given such a small likely contribution of contaminating template to gut libraries, we did not exclude gut OTUs based on information from the negative controls.

To normalize coverage we down-sampled all libraries in the OTU table (including those from both reproducibility and repeatability experiments) to 105,000 sequences each, which we deemed an optimal tradeoff between sequencing depth and retention of samples for analysis. This lead to the exclusion of four libraries from the reproducibility study: all three samples from one FW fish (bringing the total number of fish analyzed in this experiment to 35) and the PCP library from a second FW fish. We then generated count summary tables for all taxonomy levels using QIIME’s summarize_taxa.py, but downstream analyses described in this report feature phylum, class, species, or individual OTU counts (see Results). We used QIIME’s core_diversity_analyses.py to generate phylogenetic diversity metrics separately for the reproducibility and precision studies, including Faith’s Phylogenetic Diversity (Faith 1992), unweighted UniFrac (Lozupone & Knight 2005), and weighted UniFrac (Lozupone *et al.* 2007). To evaluate the potential influence of erroneous sequences on some downstream analyses, we also generated unweighted and weighted UniFrac dissimilarity matrices for the reproducibility study based on OTU picking with a sequence identity threshold of 90%. All downstream analyses were based on the respective down-sampled count tables and phylogenetic diversity metrics described above, and were conducted using version 3.3.2 of the R statistical language (R Core Team 2016).

#### Reproducibility of alpha and beta diversity metrics

We evaluated relative contributions of biological (among-fish) and technical (within-fish) variation using a repeated measures linear model framework. In particular, and following Lessels and Boag (1987), we calculated “repeatability” (reproducibility in this case) as:

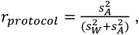

where 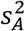 is the among-fish variance component and 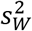 is the within-fish (among-protocol) variance component, as calculated from mean squares in an ANOVA (Lessells & Boag 1987). *r_protocol_* ranges from 0 to 1, with high values indicating increasingly small contributions of DNA isolation protocol relative to individual host contributions, which are of interest to ecologists and evolutionary biologists. A perfect reproducibility of 1.0, for example, would indicate zero within-fish variance, that is, no effect of protocol in this situation. Alternatively, a repeatability of 0.5 would be interpreted as equal variance contributions from individual and protocol. We calculated reproducibility for class richness, class evenness, species richness, species evenness, and Faith’s Phylogenetic Diversity using variance components estimated by the lmer function from the R package lme4 (Bates *et al.* 2015). For each metric we also resampled (with replacement) 35 individual fish 500 times, and used the distribution of reproducibility values from the 500 bootstrap replicates to calculate 95% confidence intervals. We also applied this approach to multivariate measures of community dissimilarity (class- and species-level Bray-Curtis dissimilarity, and weighted and unweighted UniFrac) by extracting the above variance components using the R function nested.npmanova from the BiodiversityR package (Kindt 2005). Furthermore, we calculated population-specific reproducibilities (and confidence intervals) for all of the above variables to evaluate whether reproducibility differed depending on host population. We also estimated reproducibility for weighted and unweighted UniFrac based on OTU definition exactly as described above, but using a 90% sequence identity threshold, to evaluate potential influence of sequencing error.

#### Testing effects of DNA isolation protocol, host population, and their interaction on alpha diversity, beta diversity, and relative taxon abundances

For the same five alpha diversity variables mentioned above, we evaluated significance of the fixed effects of protocol and stickleback population using mixed linear models that included the random effect of individual nested within family. Note that family is indistinguishable from tank in our design, so family and tank effects cannot be separated. We fit full and reduced models using lmer from the R package lme4 (Bates *et al.* 2015) and tested null hypotheses of no population, protocol, and population-by-protocol interaction effects on each diversity variable using likelihood ratio tests.

We also tested the influence of these factors on four measures of community dissimilarity (beta diversity) using two permutational analysis of variance (PERMANOVA) tests (Anderson 2001). First, we evaluated the effects of DNA isolation protocol, family (tank), and their interaction using the adonis2 function from the R package vegan (Oksanen 2017). Second, we evaluated the effect of population, accounting for non-independence of individuals within the same family (tank), separately for the three DNA isolation protocols using the function nested.npmanova from the BiodiversityR package (Kindt 2005). Finally, to test whether among-fish community dissimilarity was correlated between DNA isolation protocol pairs we performed Kendall’s *τ*-based Mantel tests using the mantel function from the R package vegan (Oksanen 2017).

To evaluate effects of DNA isolation and stickleback population on relative abundances of class-level and species-level OTU groups (“L3” and “L7,” respectively, from summarize_taxa.py), we fit generalized linear mixed models that included the random effect of individual nested within family. We considered only those taxonomy groups represented by at least five counts in at least nine libraries. Given the overdispersed nature of these count data, we fit Poisson-lognormal models using the glmer function from the R package lme4 (Bates *et al.* 2015) by including an observation-level effect in each model and by specifying the “poisson” family of generalized linear model. Because 16S data provide information about relative, as opposed to absolute, abundances of the organisms in each sample, it should be acknowledged that differences in OTU and taxonomic group counts among samples could reflect compositional differences in the community as opposed to organism-specific ones. We evaluated the significance of each effect for each taxonomy group using False Discovery Rate-controlled (Benjamini & Hochberg 1995) likelihood ratio tests, Akaike information criterion (AIC) and Bayesian information criterion (BIC).

#### OTU rarity and reproducibility of relative OTU abundance estimates

We measured the reproducibility of relative abundance estimates for 2278 individual OTUs that were present in one or more libraries from at least 10 of the 35 fish from our reproducibility experiment. We estimated reproducibility using the same general repeated measures framework above, except that we used the R package rptR (Stoffel *et al.* 2017) to calculate reproducibility and its 95% confidence interval for each OTU. We used the rpt function of rptR because it allows the flexibility of fitting an over-dispersed Poisson generalized linear model and implements computationally efficient CI construction by parametric bootstrapping. We characterized the relationship between OTU abundance reproducibility and average OTU abundance by fitting three logistic models: one for the point estimate, one for its 95% CI upper bound, and one for its 95% CI lower bound. The logistic model parameterization was as follows:

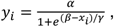

where *y* is the reproducibility of abundance, its CI upper bound, or its CI lower bound for OTU *i, x* is the logarithm to base 10 of the among-library mean abundance of OTU *i, α* is the asymptote, *β* is the inflection point in units of *x*, and *γ* is the steepness of the relationship at inflection. We fit these logistic models using the R package nls2 (Grothendieck 2013).

#### Diversity metric precision

We measured the precision with which each of several alpha and beta diversity metrics was estimated, comparing across the three DNA isolation protocols. This was made possible by sampling two guts for each protocol, and six technical replicates per gut (Fig 1B). In this case within-fish variance was attributable only to tissue subsampling and technical differences among the 6 libraries prepared identically, and not protocol differences. Because we sampled only two fish for each DNA isolation protocol, we could not effectively apply the repeatability framework used in the reproducibility study (see above), which relies on adequately sampling across-individual variation. Instead, we conducted Levene’s tests according to Sokal and Rohlf (2011) to test whether the average absolute deviation from group (individual gut) medians was significantly different among the three DNA isolation protocols. We applied this test to class- and species-level richness and evenness, and Faith’s Phylogenetic Diversity. We conducted the same type of test for class- and species-level Bray-Curtis Dissimilarity, and weighted and unweighted UniFrac, but we considered distance between observation and group (individual gut) centroid as the response variable.

## RESULTS

### Tissue subsampling and two-tiered bead beating improve gut DNA yield and integrity

We compared DNA yield and fragment size distribution between guts first homogenized with steel beads, subsampled, then treated with a second bead beating step aimed at microbial lysis, against guts handled without these modifications. By reducing tissue mass through measured, consistent subsampling, and by including the second bead beating step, we achieved higher DNA yield and integrity, and lower variance among individuals (Fig 2). Subsampled, double-beat DNA isolates contained more micrograms of DNA on average (Fig 2 A-B). For column-based DNA isolation protocols (PFP and DEP), median yield increased at least 2-fold. DNA integrity also improved with the modifications (Fig 2 C-D), especially in the case of phenol-chloroform (PCP) isolations. Because whole guts were used for the unmodified protocols, within-fish comparisons of these unmodified protocols, and testing of protocol-by-fish interactions, were not possible.

**Figure 2.**
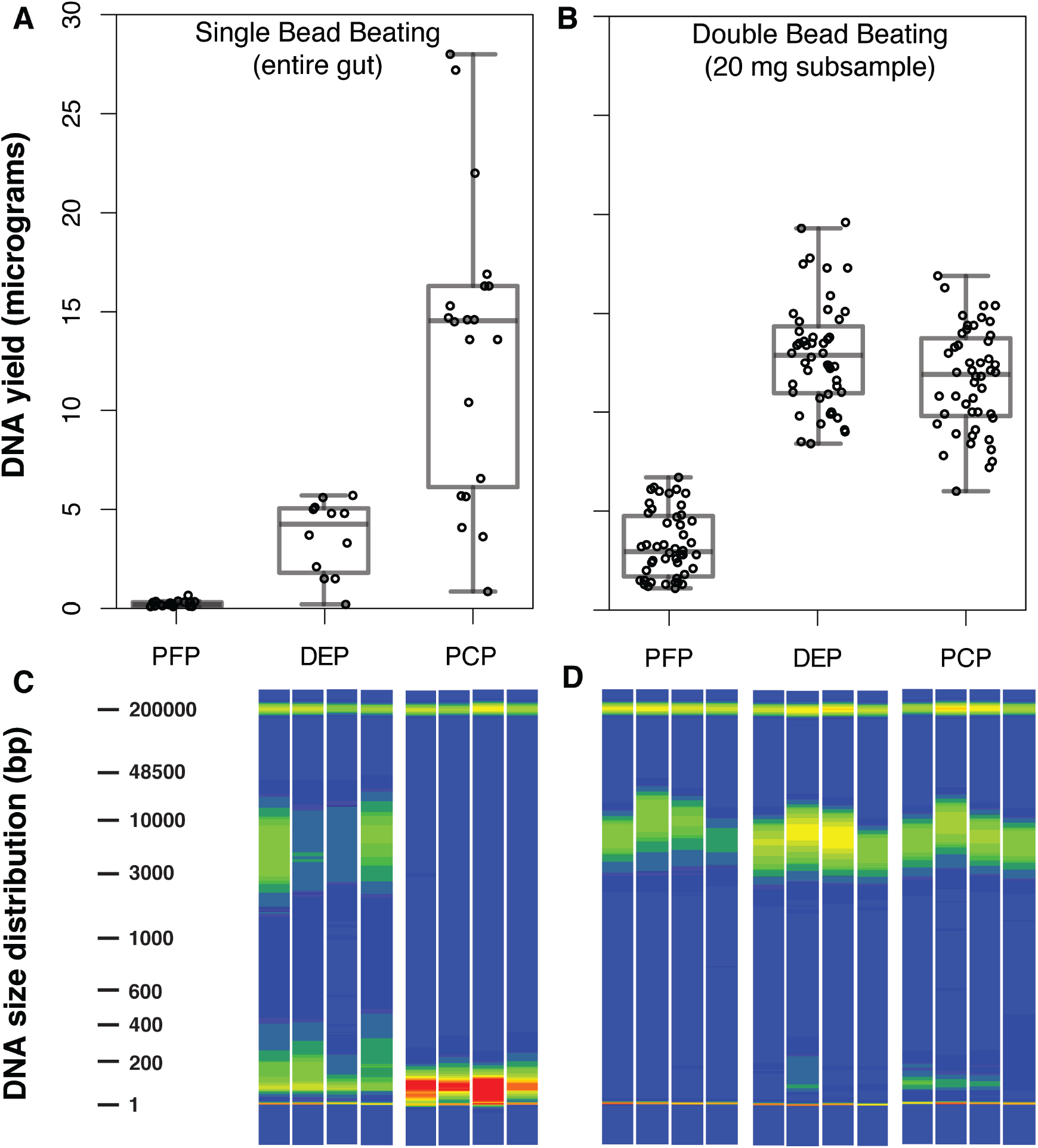
Gut homogenate subsampling and double bead beating improves DNA yield and integrity. **A**. DNA yield boxplot for 3 protocols with single beating and no gut subsampling. **B**. DNA yield boxplot for 3 protocols with double beating and subsampling. **C**. Fragment analysis traces for 2 protocols with single beating and no subsampling. **D**. Fragment analysis traces for 3 protocols with double beating and subsampling. Bands at 1 and 200,000 bp in C-D are lower and upper size standards. Fragment analysis data were unavailable for singly beat PFP samples owing to insufficient DNA quantity. PCP = Phenol-chloroform protocol, PFP = PowerFecal protocol, and DEP = DNeasy protocol.

Importantly, the among-sample variation, as estimated by the coefficient of variation for all three DNA isolation protocols, was also lower after subsampling and double bead beating (PCP: 0.557 vs 0.223; PFP: 0.610 vs. 0.509; DEP: 0.517 vs. 0.214; single vs. double respectively). The coefficient of variation for DNA yield across all singly beat, whole gut samples was 1.244, compared to 0.527 for doubly beat, subsampled guts. This difference was significant based on the asymptotic test described by Feltz and Miller (1996), which assumes a *χ*^2^- distributed test statistic (D’AD = 37.376; df = 1; *p* = 9.742e-10).

### Bacterial phyla of the stickleback gut microbiome are similar across studies and rearing environments

Rarefaction curves based on samples from 10 to 150,000 sequences per library indicated that our final down-sampling threshold of 105,000 sequences captured reasonable alpha diversity given the rate of increase with sampling effort (Fig S1A-B). Considering all 41 experimental fish (35 from the reproducibility experiment and 6 from the repeatability experiment) for which our sequence number threshold of 105,000 was reached, we recovered a mean per-individual OTU richness of 4378.122 (SEM = 291.360), which reflects OTUs summed across all libraries (each library downsampled to 105,000 sequences) per individual. At the phylum level we observed a mean richness of 28.146 (SEM = 0.786). The major constituent phyla among the fish in our experiment included Proteobacteria, Firmicutes, Chloroflexi, Bacteroidetes, and Cyanobacteria, but we observed extensive among-individual variation (Supplementary Fig S1D). Phylum-level membership was comparable between the lab-reared fish in our study and both lab-reared and wild-caught fish from Bolnick et al. (2014), with the exception of greater relative abundance of Phylum Chloroflexi in our study (Supplementary Fig S1C).

### Effects of individual hosts on microbial diversity are much larger than those of DNA isolation protocols

We evaluated the relative contributions of individual fish and DNA isolation protocol to overall variance in diversity metrics by treating the libraries from the three different protocols as repeated measurements of each fish. The contribution of protocol to variation in community composition was quite small relative to that of individual fish at class (Fig 3) and species (Supplementary Fig S2) levels, as quantified by high reproducibility estimates for all five alpha diversity metrics (Table 1). Similarly, the effect of protocol on beta diversity was weak relative to that of individual (Fig 4; Supplementary Fig S3), as reproducibility with respect to class and species Bray-Curtis dissimilarity and weighted UniFrac was high. Interestingly, reproducibility was substantially lower for unweighted UniFrac (Table 1; Fig 4). We also observed low reproducibility for unweighted, relative to weighted, UniFrac after defining OTUs based on 90% sequence identity (Table 1), which should absorb any effects of sequencing error. Confidence intervals (95% bootstrap) for reproducibility calculated separately for the two stickleback populations overlapped for all nine diversity metrics (Table 1), suggesting that reproducibility was consistent for the two different host genetic backgrounds.

**Figure 3.**
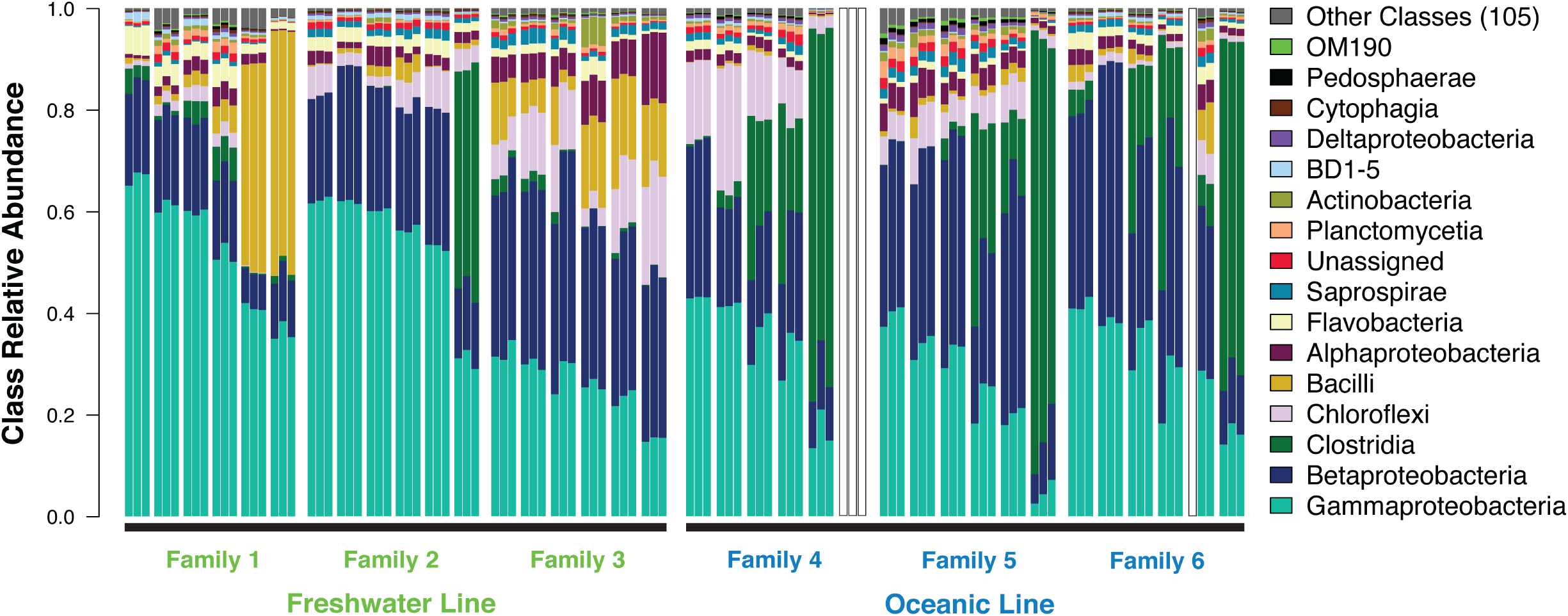
16S-based, class-level profiles of the stickleback gut microbiome vary substantially more by individual host than by DNA isolation method. Class relative abundances demonstrate substantial variation across individuals, families, and populations, but little variation among DNA isolation protocols within individuals. Each bar triplet denotes an individual fish gut, with individual bars representing PCP, PFP, and DEP DNA isolation methods, in that order. Individuals are sorted by mean Gammaproteobacteria abundance within each family. One individual from Family 4 and the PCP library from a Family 6 individual were not analyzed owing to insufficient coverage.

**Figure 4.**
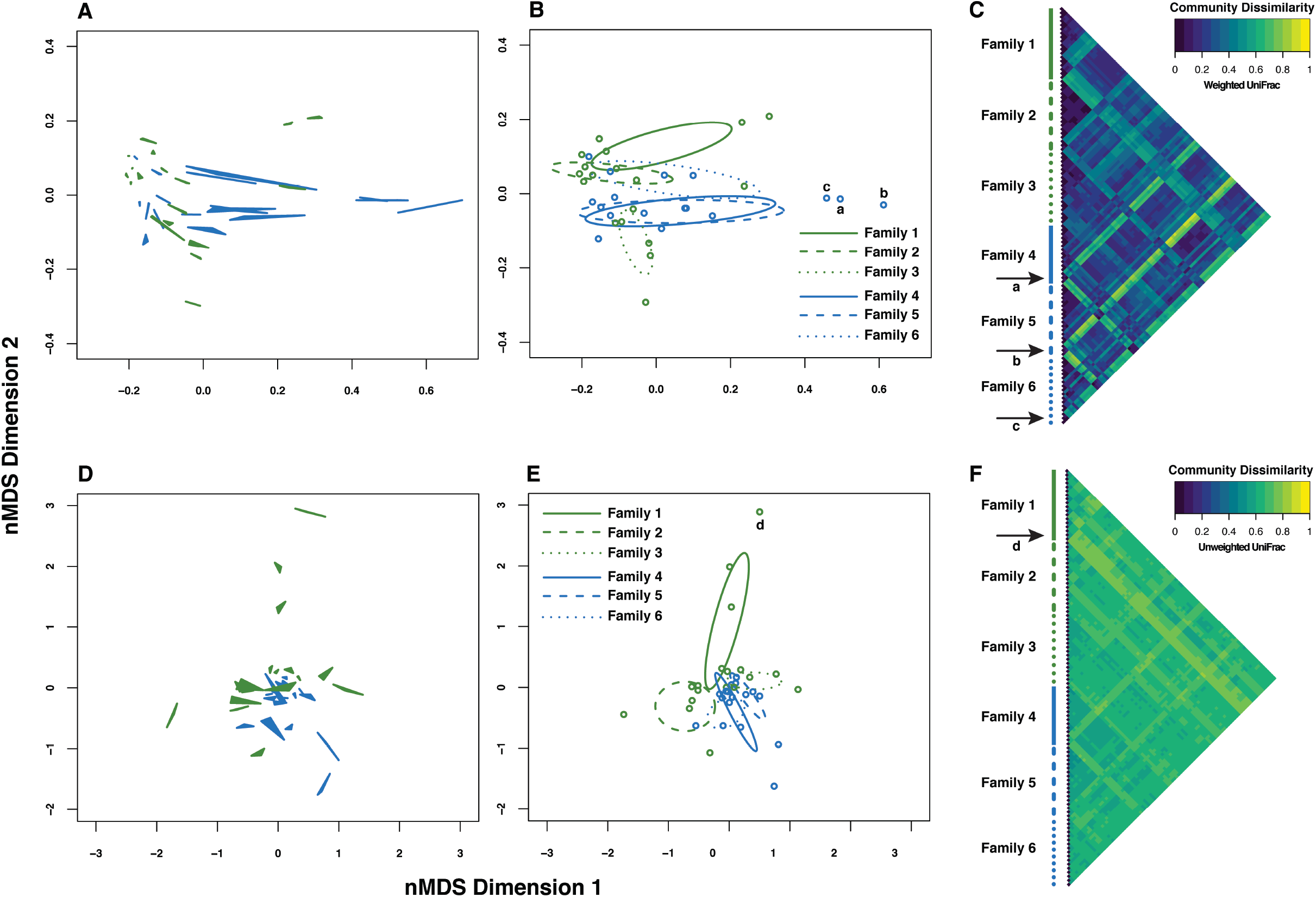
Phylogenetic dissimilarity based on 16S profiles of the stickleback gut microbiome shows a greater effect of individual, relative to DNA isolation protocol. The strength of this pattern varies, depending on whether weighted (**A-C**) or unweighted (**D-F**) UniFrac is applied. **A** and **D**. Non-metric multidimensional scaling (nMDS) ordinations from weighted and unweighted UniFrac, showing the three DNA isolation protocols from each individual connected as filled triangles. **B** and **E**. The same ordinations, but with individuals plotted as the centroid of each triplet from **A** and **D**, and with 95% confidence ellipses drawn separately for each family. **C** and **F**. Pairwise dissimilarity matrix heatmaps representing all libraries. The library order is the same as in Fig 3. Green ordination symbols represent the freshwater stickleback line, and blue symbols represent the oceanic line. Individual fish labeled by lowercase letters and corresponding arrows point to outliers in community space.

**Table 1.**
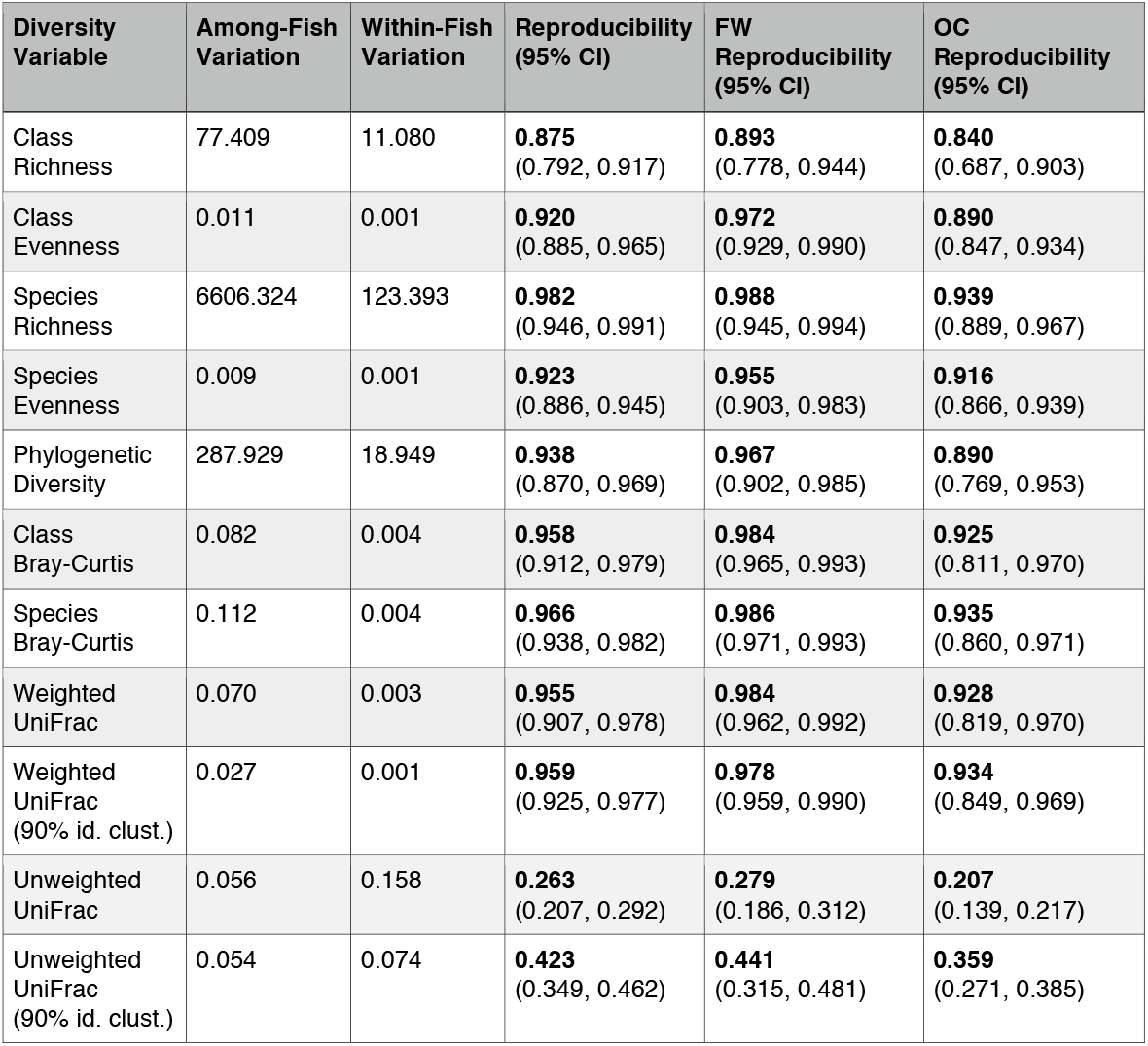
Among-fish variation greatly exceeds within-fish (among-protocol) variation for several diversity metrics, as indicated by high reproducibility estimates. Shown are variance component and reproducibility estimates for all individuals, and separate reproducibility estimates for freshwater (FW) and oceanic (OC) stickleback populations, respectively, along with bootstrap 95% confidence intervals.

| **Diversity Variable** | **Among-Fish Variation** | **Within-Fish Variation** | **Reproducibility (95% CI)** | **FW Reproducibility (95% CI)** | **OC Reproducibility (95% CI)** |
| --- | --- | --- | --- | --- | --- |
| Class Richness | 77.409 | 11.080 | **0.875**<br>(0.792, 0.917) | **0.893**<br>(0.778, 0.944) | **0.840**<br>(0.687, 0.903) |
| Class Evenness | 0.011 | 0.001 | **0.920**<br>(0.885, 0.965) | **0.972**<br>(0.929, 0.990) | **0.890**<br>(0.847, 0.934) |
| Species Richness | 6606.324 | 123.393 | **0.982**<br>(0.946, 0.991) | **0.988**<br>(0.945, 0.994) | **0.939**<br>(0.889, 0.967) |
| Species Evenness | 0.009 | 0.001 | **0.923**<br>(0.886, 0.945) | **0.955**<br>(0.903, 0.983) | **0.916**<br>(0.866, 0.939) |
| Phylogenetic Diversity | 287.929 | 18.949 | **0.938**<br>(0.870, 0.969) | **0.967**<br>(0.902, 0.985) | **0.890**<br>(0.769, 0.953) |
| Class Bray-Curtis | 0.082 | 0.004 | **0.958**<br>(0.912, 0.979) | **0.984**<br>(0.965, 0.993) | **0.925**<br>(0.811, 0.970) |
| Species Bray-Curtis | 0.112 | 0.004 | **0.966**<br>(0.938, 0.982) | **0.986**<br>(0.971, 0.993) | **0.935**<br>(0.860, 0.971) |
| Weighted UniFrac | 0.070 | 0.003 | **0.955**<br>(0.907, 0.978) | **0.984**<br>(0.962, 0.992) | **0.928**<br>(0.819, 0.970) |
| Weighted UniFrac (90% id. clust.) | 0.027 | 0.001 | **0.959**<br>(0.925, 0.977) | **0.978**<br>(0.959, 0.990) | **0.934**<br>(0.849, 0.969) |
| Unweighted UniFrac | 0.056 | 0.158 | **0.263**<br>(0.207, 0.292) | **0.279**<br>(0.186, 0.312) | **0.207**<br>(0.139, 0.217) |
| Unweighted UniFrac (90% id. clust.) | 0.054 | 0.074 | **0.423**<br>(0.349, 0.462) | **0.441**<br>(0.315, 0.481) | **0.359**<br>(0.271, 0.385) |

Although the overall variance in diversity metrics explained by differences in DNA isolation protocol was small relative to that explained by among-individual differences, we detected a significant effect of protocol for some measures via likelihood ratio tests comparing full and reduced linear mixed models (Table 2; Supplementary Figs 4-5). For example, total variation in class richness was explained significantly better by a model including DNA isolation protocol, relative to one excluding the term. This effect size was small, however (Supplementary Fig S4A). Libraries from DNA isolated using DNeasy (DEP) yielded a modest increase in mean class richness from 52.663 to 54.892, with respect to PowerFecal (PFP), and 53.528 to 54.892 with respect to Phenol-Chloroform (PCP). We observed a similar trend for species richness and Faith’s Phylogenetic Diversity (Table 2; Supplementary Fig S5A,C).

**Figure 5.**
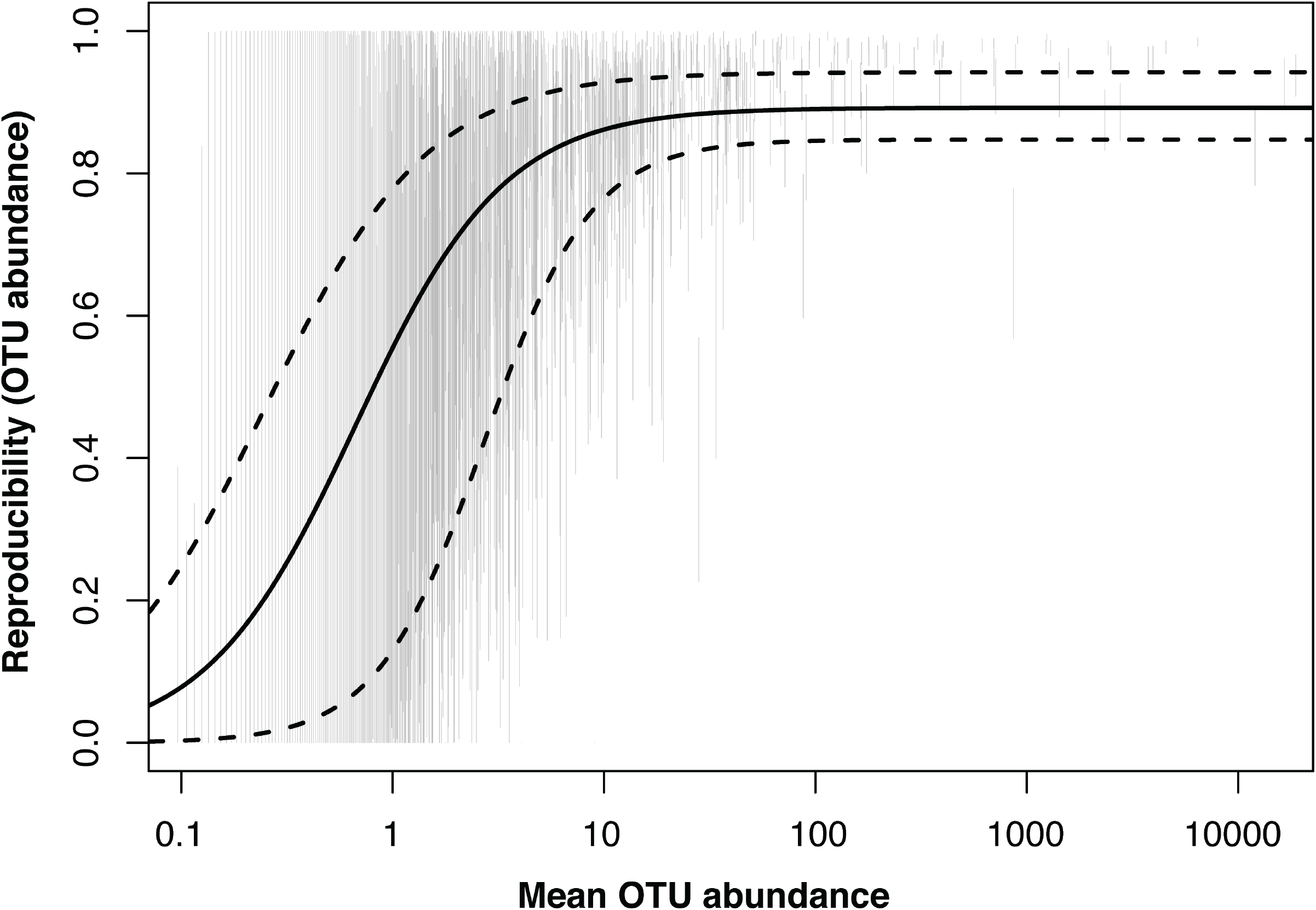
Repeatability of OTU quantification across DNA isolation protocols increases nonlinearly with log_10_-transformed mean relative OTU abundance. Vertical gray lines represent 95% CIs for repeatability estimates of 2278 OTUs observed in at least 10 of 35 experimental fish. The solid line represents predicted repeatability values from a logistic model fit to the data. Dashed lines represent predicted upper and lower bound CI values for repeatability, also from logistic models fit to the data.

**Table 2.**
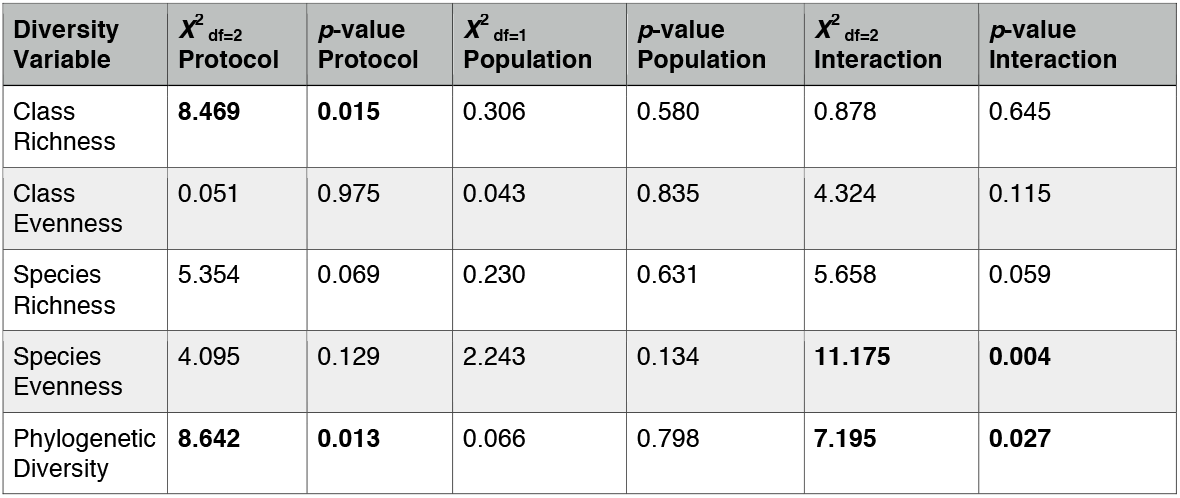
Likelihood Ratio Test (LRT) test statistics with degrees of freedom (df) and *p*-values for tests of effects of DNA isolation protocol, stickleback population, and interaction between the two. LRTs were conducted by comparing linear mixed models either including or excluding these fixed effects, plus random effects of fish and fish nested within family (see Methods). Tests with *p*-values < 0.05 are in bold.

Beta diversity (as measured by class- and species-level Bray-Curtis dissimilarity, unweighted Unifrac, and weighted Unifrac) was significantly influenced by DNA isolation protocol, on average, in all cases (*p* < 0.001; see Supplementary Sheet S1A for factorial PERMANOVA hypothesis test statistics). The effect sizes were once again quite small in the context of among-individual variation, as reflected in nMDS ordinations and pairwise library dissimilarity distributions (Fig 4A,C,D,F; Supplementary Fig S3A,C,D,F).

Relative abundances of individual taxon groups were in some cases affected by DNA isolation protocol, based on comparison of lognormal Poisson generalized linear models. For 20 class-level and 89 species-level OTUs, the model including protocol was a better fit than the model excluding it based on AIC, BIC, and the FDR-controlled likelihood ratio test (Supplementary Sheet S1 B-C). For instance, the class-level groups Actinobacteria, BD1-5 (“Gracilibacteria”), and Thermomicrobia tended to vary in abundance among DNA isolation protocols within fish consistently (Supplementary Fig S6A-C), albeit with quite small effect sizes. The mean downsampled read count for Actinobacteria, for example, was 1. 317 times higher for PCP relative to PFP methods. At the species level, taxonomy groups including *Agromyces spp.*, an unassigned species from Family Rodobacteraceae, and *Tsukamurella spp.* were among the most likely taxa affected by DNA isolation protocol (Supplementary Fig S7A-C), also to a minor degree.

We wanted to evaluate the potential for DNA isolation protocol differences to influence the ability to consistently measure among-host differences in relative OTU abundances, and whether this ability varied as a consequence of the scarcity of a given OTU. We found that reproducibility was indeed positively associated with average log_10_ OTU abundance, based on a fitted logistic model (Fig 5). The slope at inflection (*γ*) was significantly greater than zero (*γ* = 0.352; std. err. = 0.015; *t* = 23.977; *p* < 0.0001). A similar relationship was observed for upper and lower 95% CI bounds on reproducibility (Fig 5). We noted a similar pattern when measuring OTU rarity in a different way: the number of individual hosts in which the OTU was detected. Again we observed that reproducibility of relative OTU abundance estimates was higher for common OTUs (Supplementary Fig S8A), and that the lower bound for reproducibility in our sample of hosts increased substantially when the OTU was present in at least 32 of 35 fish (Supplementary Fig S8B).

### Effects of stickleback population on microbial diversity estimates are subtle and in limited cases contingent on DNA isolation protocol

We did not detect a statistically significant effect of host population (stickleback line) on any of the five alpha diversity metrics using likelihood ratio tests comparing nested full and reduced linear mixed models (Table 2), although class richness, class evenness, and species richness trended towards higher values in the oceanic relative to the freshwater population (Supplementary Figs 4-5). We did detect a statistically significant interaction between stickleback population and DNA isolation protocol for species-level and phylogenetic alpha diversity metrics (Table 2). This implies that the effect of isolation protocol may differ depending on biological context, but these effect sizes were also quite small (Supplementary Fig S5B-C). For example, the maximum species evenness difference between protocol-population combinations was 0.070 and the maximum phylogenetic diversity difference was 5.370.

Beta diversity, as measured by class- and species-level Bray-Curtis dissimilarity, unweighted Unifrac, and weighted Unifrac, was not significantly influenced by host population after accounting for the nested nature of the data introduced by family structure (Fig 4B,C,E,F; see Supplementary Sheet S1A for protocol-specific nested PERMANOVA hypothesis test statistics). Family itself was a significant determinant of community dissimilarity for all four metrics (Fig 4B,C,E,F; see Supplementary Sheet S1A for factorial PERMANOVA hypothesis test statistics), although it should be noted that family was confounded by tank in our design. We did not detect a statistically significant interaction between host family and DNA isolation protocol for any of the four dissimilarity metrics assessed (Supplementary Sheet S1A). Finally, among-fish community dissimilarity ranks were highly correlated between DNA isolation protocols (Supplementary Sheet S1D), suggesting consistency with respect to between-individual beta diversity, especially for PFP and DEP protocols.

We also evaluated whether relative abundances for taxonomic groups (class- and species-level) might be affected by host population, based on comparison of lognormal Poisson generalized linear models accounting for family nestedness. Although no likelihood ratio tests were statistically significant after controlling the FDR at 0.1, three class-level and 25 species-level groups showed evidence for a population effect by virtue of a delta AIC > 2, a delta BIC > 0, and an uncorrected LRT *p*-value < 0.05 (Supplementary Sheet S1B-C). Class-level groups BD-7, an unassigned class from Bacteroidetes, and Clostridia were all three enriched in abundance in the oceanic relative to the freshwater population (Supplementary Fig 6D-F). The effect size of population on the Clostridia group abundance, for example, was quite large. The mean oceanic Clostridia count was 23259.559 (SEM = 3880.291), whereas the mean freshwater Clostridia count was 3867.704 (SEM = 1417.775). Three species-level groups with especially strong tendencies toward host population differences were the oceanic-enriched *Sphingobacterium multivorum* group, the freshwater-enriched *Plesiomonas shigelloides* group, and an oceanic-enriched, unassigned group from Family *Clostridiaceae* (Supplementary Fig S7D-F).

We detected a statistically significant interaction between host population (accounting for family) and DNA isolation protocol for six class-level and 25 species-level groups (Supplementary Sheet S1B-C), based on a delta AIC > 2, a delta BIC > 0, and a LRT FDR controlled at 0.10. However, effect sizes for this interaction type were again relatively small, as demonstrated by the abundance of a *Sphingobacterium multivorum* group across population-protocol combinations (Supplementary Fig S7D). In this case, the mean population difference in *S. multivorum* count was highest for DEP (4.497), followed by 1.688 and 2.912 for PCP and PFP, respectively.

### Precision of gut microbiome diversity measurements is high and similar across three DNA isolation protocols

We performed repeated measurements of individual stickleback gut microbiomes obtained from replicate aliquots from whole gut homogenates and using a single DNA isolation protocol per gut (Fig 1B). 16S data generated from the same fish were extremely similar in taxonomic composition, relative to among-fish comparisons (Supplementary Figs S9A and S10A). We analyzed within-individual variation based on the six gut subsamples per fish and found no significant effect of protocol on precision for five alpha diversity metrics (Supplementary Sheet S1E), including class-level richness and evenness (Supplementary Fig S9), species richness and evenness (Supplementary Fig S10A-C), and phylogenetic diversity (Supplementary Fig S10E). Similarly, we found no evidence for a significant effect of protocol on precision with respect to beta diversity (See Supplementary Sheet S1D), including class- and species-level Bray-Curtis dissimilarity and weighted and unweighted UniFrac (Fig 6; Supplementary Fig S11). Furthermore, and consistent with our across-protocol reproducibility analysis, the average degree of within-fish, relative to among-fish, dispersion was especially low for unweighted UniFrac (Fig 6).

**Figure 6.**
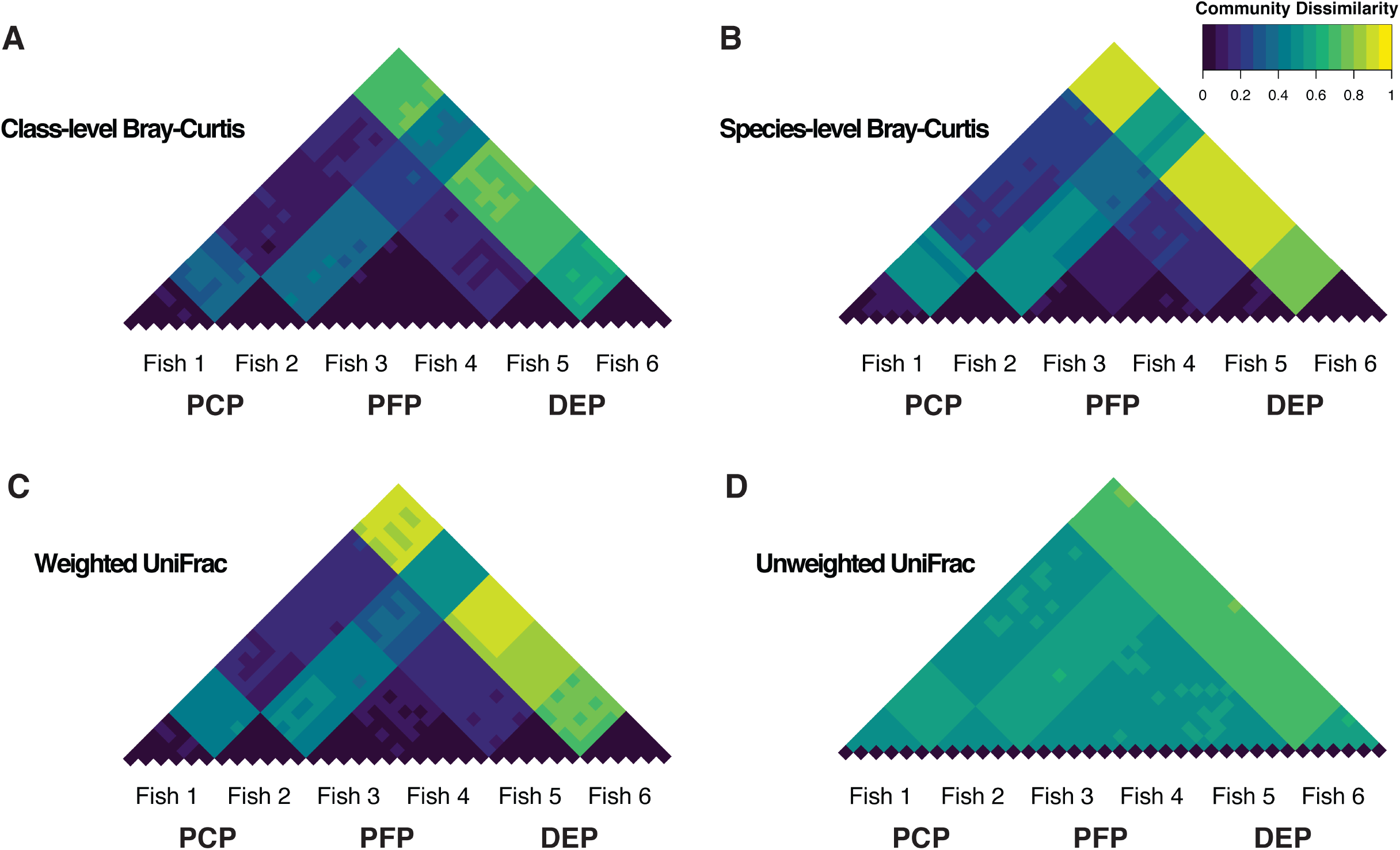
Precision of beta diversity measurements is consistently high among DNA isolation methods, but the relative magnitude of within- and among-fish community dissimilarity depends on the dissimilarity metric. Pairwise dissimilarity matrix heatmaps for the precision experiment, including class- and species-level Bray-Curtis (**A-B**), and weighted and unweighted UniFrac (C-D), illustrate low within-fish dissimilarity and higher among-fish dissimilarity. This pattern, however, is less extreme for unweighted UniFrac.

## DISCUSSION

One surprising insight from the recent characterization of microbiomes using high-throughput sequencing has been the extensive diversity among individual hosts of the same species (Adair *et al.* 2018; Human Microbiome Project 2012; Kueneman *et al.* 2014; Sullam *et al.* 2015). The fundamental sources of this inter-individual variation remain an active area of research. Rather than assuming that individual variation is vastly larger than technical variation, and therefore insignificant, we and others (Knight *et al.* 2018; Poussin *et al.* 2018; Sinha *et al.* 2017) argue that the relative magnitude of biological and technical variance components should be measured using strong experimental design. This is particularly relevant in the context of new and rapidly changing technologies for quantifying microbial diversity. Indeed, variation in microbial diversity metrics may be heavily influenced by technical factors in some cases, especially when molecular protocols are vastly different or suboptimal. For instance, extremely low yields of microbial DNA exacerbate the influence of contaminant species (Salter *et al.* 2014), which could negatively impact biological inference. Furthermore, inferences based on some diversity metrics might be more susceptible to technical variation, particularly those metrics more heavily influenced by sequences from rare species whose abundance estimates may be more subject to sampling error.

In our experience, suspicions and intuition about the severe importance or unimportance of technical variation for 16S-based microbial ecology inference have been extensively discussed. However, these effects have not been precisely quantified and evaluated outside the purview of “mock communities” (Salter *et al.* 2014; Yuan *et al.* 2012) or differences among research groups working on large-scale, collaborative efforts to understand the human microbiome (Sinha *et al.* 2017). While these studies and several others, for example (Kashinskaya *et al.* 2017; Lawrence *et al.* 2015; Raju *et al.* 2018; Videvall *et al.* 2017; Wagner Mackenzie *et al.* 2015), have been useful in identifying *potential* sources of technical variation that may or may not restrict or bias biological inferences based on among-individual variation, insufficient biological replication and/or inability to isolate specific technical factors have limited their scope of inference. For example, authors from the Microbiome Quality Control Project point out that their carefully designed study was “unable to assign significance to any specific fixed effects (i.e., individual protocol variables), since in the small MBQC-base these were in large part confounded with individual handling and bioinformatics laboratories” (Sinha *et al.* 2017).

No published study, to our knowledge, has sampled enough individual hosts, in combination with the controlled assignment of technical factor levels, to effectively quantify reproducibility (and its uncertainty) in a biologically relevant context. We wanted to fill this important void with a well-replicated comparison involving one potential source of technical variation - DNA isolation protocol - and individual-level variation in the ecological and evolutionary context of stickleback host genetic differences.

One of our significant findings is that the earliest steps in sample handling are critical for improving downstream results. We found the process of dual bead beating with tissue homogenate subsampling essential to the quantity and quality of DNA isolated from adult threespine stickleback guts. Without this process, both lower yields with higher variance and more fragmented DNA were certain. The PowerFecal column-based isolation protocol suffered most severely from a lack of double bead beating and subsampling, perhaps owing to an overloading of the column. These results are significant, as low DNA yields are known to amplify any effects of contamination (Salter *et al.* 2014). Decreasing among-sample variance in DNA attributes such as quantity, therefore, should reduce non-biological variation among 16S-based microbial profiles. In principle this will increase the power of statistical analyses, thereby reducing cost in the number of biological samples needed. In the specific case of our study, these modifications were absolutely essential to establishing a reasonable comparison of DNA isolation protocols. Before embarking on 16S sequencing for a large study, we recommend similar subsampling and optimization for large sample types, or sample types that haven’t yet been tested with commercial kits. We also recommend that DNA yield and quality distributions for at least a random subset of samples be reported in published 16S and metagenomics studies.

Our results confirm that among-individual differences in stickleback gut communities are extensive (Bolnick *et al.* 2014), consistent with work on the gut microbiome of human (Human Microbiome Project 2012) and other hosts (Kreisinger *et al.* 2014; Sullam *et al.* 2015). Although mean relative abundances of phyla in our experimental fish were qualitatively similar to those of other stickleback populations and environments (Bolnick *et al.* 2014), we found substantial variation in community composition even among male full siblings housed in the same tank. This is significant, as most ecological, evolutionary, and biomedical studies of host-associated microbes rely on an understanding of among-host differences in the microbiome. However, previous studies have not satisfactorily quantified the extent to which observed individual differences might be due to technical variation introduced by factors such as DNA isolation protocol. We measured the reproducibility (across three DNA isolation protocols) for a number of commonly used microbial diversity metrics. We found that alpha and beta diversity measurements of the stickleback gut microbiome were very reproducible, despite having applied three fundamentally different DNA isolation protocols. As stated previously, this high reproducibility is predicated upon the proper initial treatment of tissue through double beating and subsampling.

Interestingly, we observed an exception to high reproducibility in the case of unweighted UniFrac, although 95% CI lower bounds were still above zero. Because unweighted UniFrac does not account for differences in 16S sequence abundance, it magnifies the effect of rare sequences (Lozupone *et al.* 2007). To evaluate whether this property was due entirely to rare, artificial OTUs originating from sequencing error (Callahan *et al.* 2016, Amir *et al.* 2018), we re-analyzed beta diversity reproducibility for unweighted and weighted UniFrac using more inclusive OTU clustering based on 90% (as opposed to 97%) sequence identity (see Table 1), and found a reduced but still substantial difference in reproducibility for these two metrics.

Our data do not suggest that unweighted UniFrac-based measures of beta diversity are not reproducible in the general sense, but rather in the case of our study they were lower compared to metrics that take abundance into account.

The interpretation of reproducibility is contingent on the level of variation researchers wish to understand, and our objective was to study reproducibility across DNA isolation protocols (and repeatability across tissue subsamples) with respect to *among-individual* variation. Unweighted UniFrac is known to be especially sensitive to sampling bias (Lozupone et al. 2011). One recent study showed that unweighted UniFrac applied to resampled sequences from the same HMP tongue dorsum libraries projected large within-relative to among-library variation, when this should be very low (Wong *et al.* 2016). This insight, along with our current study, suggests that sampling bias associated with rare sequences disproportionately affects the potential to explain among-individual variance with unweighted UniFrac as compared to other metrics, an important consideration that researchers should make when interpreting microbiome data, especially in light of differences between similar individual hosts.

Although reproducibility among DNA isolation methods was extremely high, we observed statistically significant effects of DNA isolation protocol on some measurements of the microbiota, including class richness, Faith’s Phylogenetic Diversity, and relative abundance for at least 20 classes. It should be noted, however, that the sizes of this effect were rather small (see Results), and our experimental design was well powered to detect the effects owing to within-individual comparisons. Nevertheless, if researchers are interested in specific microbial lineages for a particular study, they should be aware that DNA protocols may indeed influence abundance estimates for individual taxa.

We also examined the relationship between average OTU abundance and among-protocol reproducibility. Based on our stickleback data, a very clear, sigmoidal relationship suggested that reproducibility was indeed lowest, on average, for rare OTUs, and that it improved substantially up to a mean OTU count of 10. The nature of this function will almost certainly vary among systems, and among sequencing depths (recall that these data were downsampled to 105,000 reads per library), but it provides a general reference for those interested in the reliable measurement of rare taxa with 16S sequencing. Even with high biological replication and relatively deep sampling, low repeatability for some organisms may be unavoidable. This pattern is fundamentally related to the reduced reproducibility we observed for unweighted UniFrac, in that increased sampling error for rare sequences makes among-individual comparisons less tractable.

We designed a second, small experiment to compare precision of 16S-based community measurements (six gut subsamples per fish) among the three DNA isolation protocols. We observed no significant difference in precision for richness, evenness, phylogenetic diversity, or beta dispersion, among DNA isolation protocols. It should be noted, however, that our sample of individual fish per protocol was limited (just two), and that among-fish variation in microbial community structure was extensive (Supplementary Fig S10A). As a result, our power to detect among-protocol differences in precision was limited.

With added confidence that optimized DNA isolation protocols contribute minimally to among-library variation, we then explored whether several factors might explain individual host differences in the stickleback gut microbiome, with a special interest in host genetic background. Recent studies of animal hosts, mostly featuring mammals, have reported a stronger influence of environmental variables relative to host genetic variation on gut microbiome variation (Bledsoe *et al.* 2018; Hildebrand *et al.* 2013; Rothschild *et al.* 2018). In our study host family significantly explained beta diversity among individuals, but family effects were confounded by those of tank. Although all tanks in our study shared a common water system, the immediate tank environment is likely to influence host-associated microbes. Future studies should address host genetic effects like those at the family level by raising individuals related to different degrees in replicated common garden experiments informed by traditional quantitative genetics principles.

The population of origin was not a statistically significant factor for most of the gut microbiome traits we measured, however individual species-level groups such as those associated with *Sphingobacterium multivorum* and *Plesiomonas shigelloides* showed strong evidence for an association with stickleback line. Taxonomic groups from Phylum Firmicutes (namely the family Clostridiaceae and the genus *Turicibacter*) also differed in abundance between the two stickleback lines. Host-genotype influences on Firmicutes appear to be common in mammals (Goodrich *et al.* 2016), and *Turicibacter* has been shown to be heritable in both humans and mice (Benson *et al.* 2010; Org *et al.* 2015). While a subtle effect of host genotype on the stickleback gut microbiome is consistent with the aforementioned insights from animal hosts, it should be interpreted with some caution, as the nested nature of our design, and having sampled only three families per population, limited our statistical power to test population-level hypotheses. Future studies that experimentally control environmental effects carefully and sample more genetic variation at the population level (e.g. genome-wide association studies and large-scale common garden experiments) should provide the power to confirm these still largely untested contributions to among-individual microbiome variation. Notably, we detected minimal evidence for statistical interaction between host population and DNA isolation protocol. Again, although some of these tests were statistically significant, the associated effect sizes were rather small (Figs S5B-C; S7D). The mechanistic causes of these subtle interactions are unknown, but it is possible that inorganic or organic compounds in the guts differing in concentration between freshwater and oceanic stickleback could co-purify with DNA and affect downstream steps in library construction such as PCR, in a manner specific to DNA from some microbial lineages.

Our current study revealed high reproducibility across the three protocols we tested, and minimal concern that choice of DNA isolation protocol interacts with biological factors of interest. In our experience the DNeasy Protocol (DEP) required the shortest handling time, so we have adopted it in current studies of the stickleback gut microbiome. Negligible influence from these technical factors may not be the case for other biological systems or sample types, however, so we strongly encourage other researchers to design their studies in ways similar to those presented here in order to properly measure and minimize sources of technical variance. The payoff in limiting technical variation is potentially large in terms of cost of reagents, time, and animal resources, especially when true biological signal is subtle. This concept is, of course, easily extended beyond 16S data sets to RNA-Seq and other high-throughput sequencing data. In summary, the complexity of communities and the sampling process can affect reproducibility and repeatability, but as we show, not always to a great extent.

The magnitude of these effects depends on the biology of the system at hand and the diversity metric in question. Without properly quantifying relevant technical and biological variance components of sequencing-based microbial diversity metrics, however, it is impossible for our research community to move forward with confidence in addressing core questions about host-microbe interactions and microbial ecology in general.

## Supporting information

## ACKNOWLEDGEMENTS

We thank K. Milligan-Myhre and Z. Stephens for important discussions that helped motivate this work. H. Tavalire provided valuable recommendations regarding the OTU rarity analysis. K. Guillemin, R. Parthasarathy, B. Bohannan, J. Eisen, and many members of the UO META Center for Systems Biology provided helpful advice and feedback throughout the project. We are also grateful to M. Weitzman and D. Turnbull from the UO GC3F for critical assistance with library preparation and sequencing. This work was funded by National Institutes of Health grants P50GM098911 (to WAC, K. Guillemin, et al.) and RR032670 (to WAC), and Oregon Research Excellence Funds (to WAC). EAB was supported by a National Institutes of Health NRSA fellowship (F32GM122419).

## CONFLICTS OF INTEREST

The authors declare no conflicts of interest.

## Data Accessibility

All 16S amplicon sequencing data associated with this study are available via the NCBI Sequencing Read Archive (SRA), under BioProject PRJNA503913.

## Author Contributions

CMS, MC, SB, and WAC conceived and designed the study. CMS and MC collected the data, and CMS performed the analyses. CMS, MC and EAB wrote an initial draft of the manuscript, and all authors contributed to subsequent versions.

## SUPPLEMENTAL INFORMATION

### Supplementary Figure Legends S1-S11

**Supplementary Figure S1.**
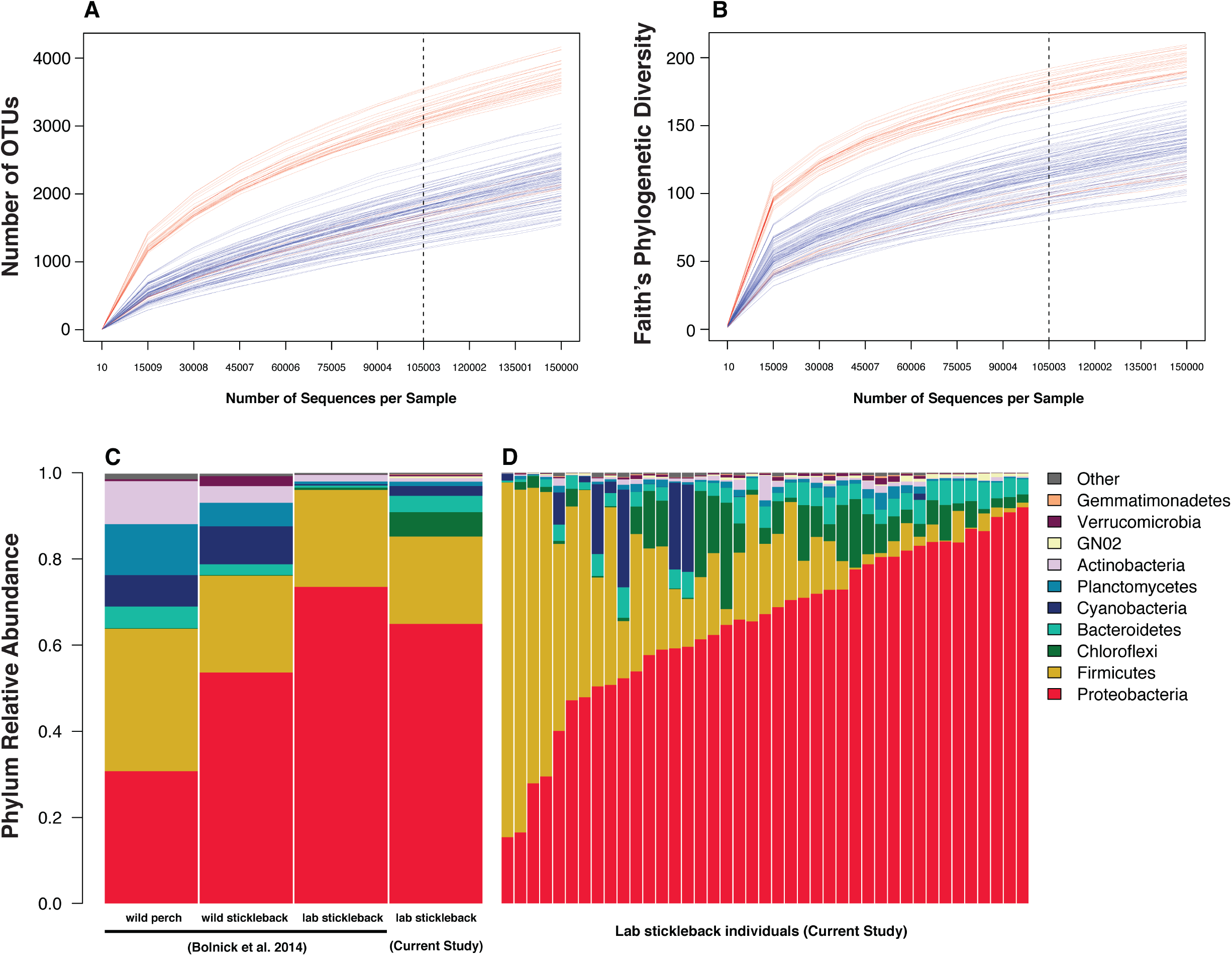
Alpha diversity rate of increase with sequencing effort decreases before the 105,000 read depth threshold. **A-B**. Library-wise rarefaction curves for OTU richness and phylogenetic diversity, respectively. The dashed lines indicate the 105,000 threshold used for all downstream analyses. Blue lines indicate libraries from the reproducibility experiment, whereas red lines indicate libraries from the repeatability experiment. On average, alpha diversity was higher for the six fish from the reproducibility experiment relative to the 36 fish from the repeatability experiment. Microbial community diversity at the phylum level in the adult threespine stickleback gut is similar across studies and rearing environments. **C**. Across-individual mean relative phylum abundances for perch, wild-caught stickleback, and lab stickleback from Bolnick et al. (2014), and lab stickleback from our current study. **D**. Individual fish phylum abundances (averaged across DNA isolation protocols) for the 41 stickleback analyzed in both experiments from the current study.

**Supplementary Figure S2.**
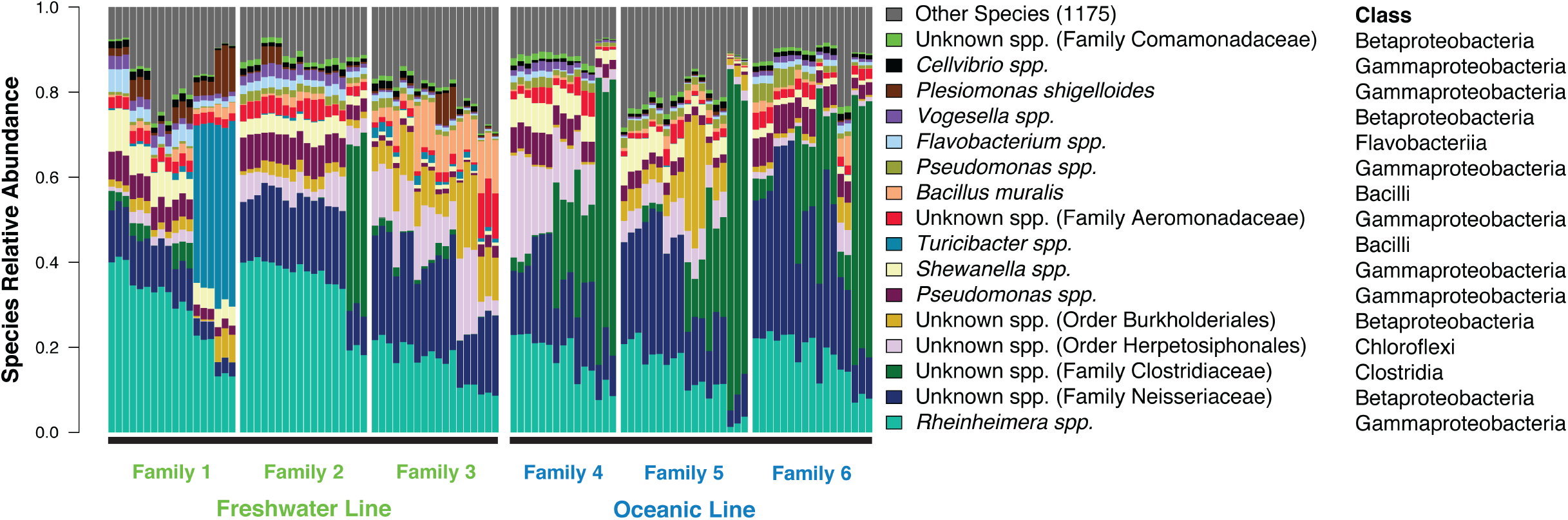
16S-based, species-level profiles of the stickleback gut microbiome vary substantially more by individual host than by DNA isolation method. Species relative abundances demonstrate substantial variation across individuals, families, and populations, but little variation among DNA isolation protocols within individuals. Adjacent bars of three denote individual fish guts, with individual bars representing PCP, PFP, and DEP DNA isolation methods, in that order. Individuals are sorted by mean *Rheinheimera spp.* abundance within each family. One individual from Family 4 and the PCP library from a Family 6 individual were not analyzed owing to insufficient coverage.

**Supplementary Figure S3.**
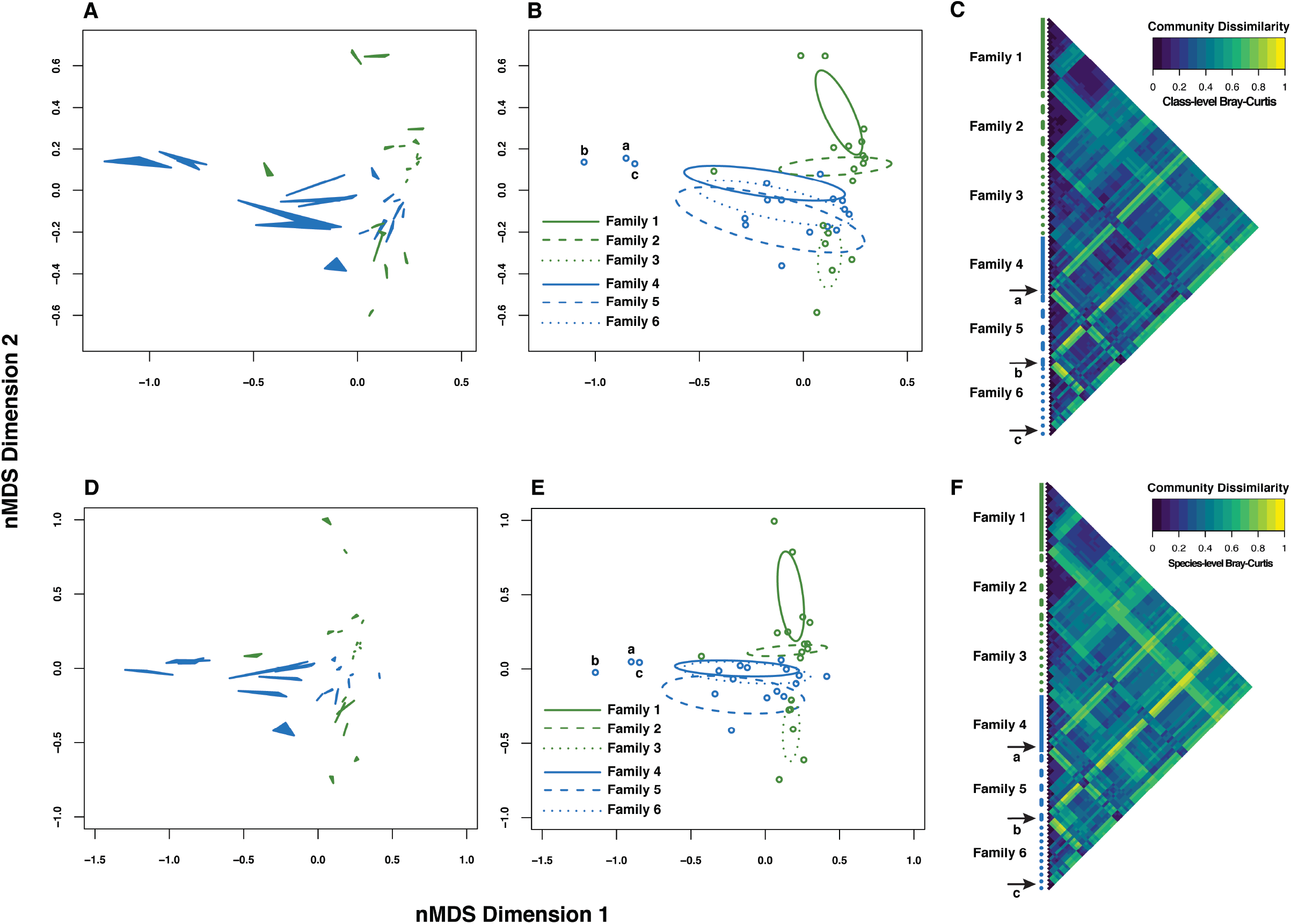
Bray-Curtis dissimilarity based on 16S profiles of the stickleback gut microbiome shows a greater effect of individual, relative to DNA isolation protocol. **A-C**. Bray-Curtis dissimilarity based on class abundances. **D-F**. Bray-Curtis dissimilarity based on species abundances. **A** and **D**. Non-metric multidimensional scaling (nMDS) ordinations from Bray-Curtis dissimilarity, showing the three DNA isolation protocols from each individual connected as filled triangles. **B** and **E**. The same ordinations, but with individuals plotted as the centroid of each triplet from **A** and **D**, and with 95% confidence ellipses drawn separately for each family. **C** and **F**. Pairwise dissimilarity matrix heatmaps representing all libraries. The library order is the same as in Fig 3. Green ordination symbols represent the freshwater stickleback line, and blue symbols represent the oceanic line. Individual fish labeled by lowercase letters and corresponding arrows point to outliers in community space.

**Supplementary Figure S4.**
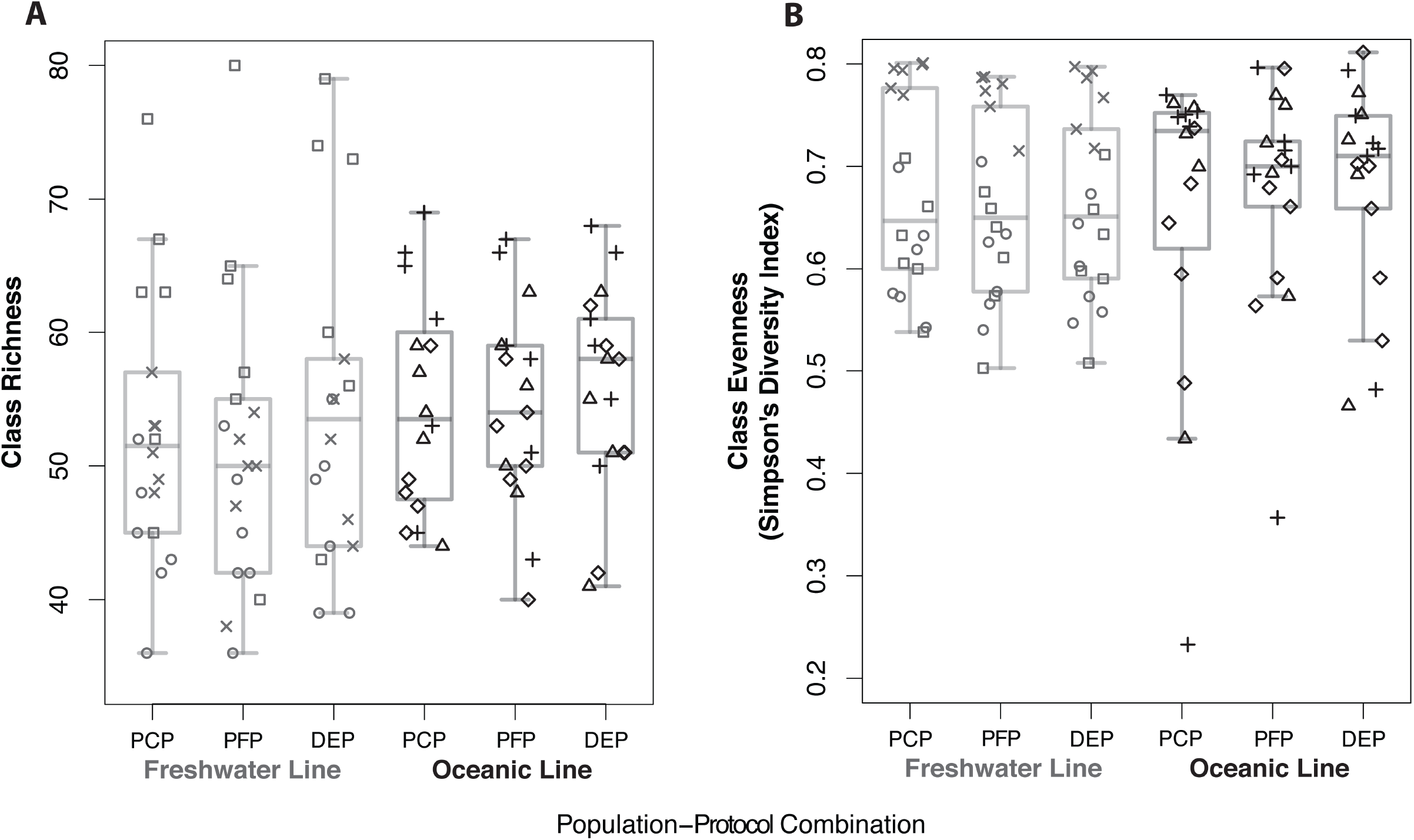
DNA isolation protocol and host population do not strongly influence class richness (**A**) or evenness (**B**). Boxplots expressing class and evenness distributions for the six population-protocol combinations are overplotted with points representing individual libraries, which are coded by stickleback family. Family 1 = square; Family 2 = circle, Family 3 = “X”; Family 4 = triangle; Family 5 = “+”; Family 6 = diamond.

**Supplementary Figure S5.**
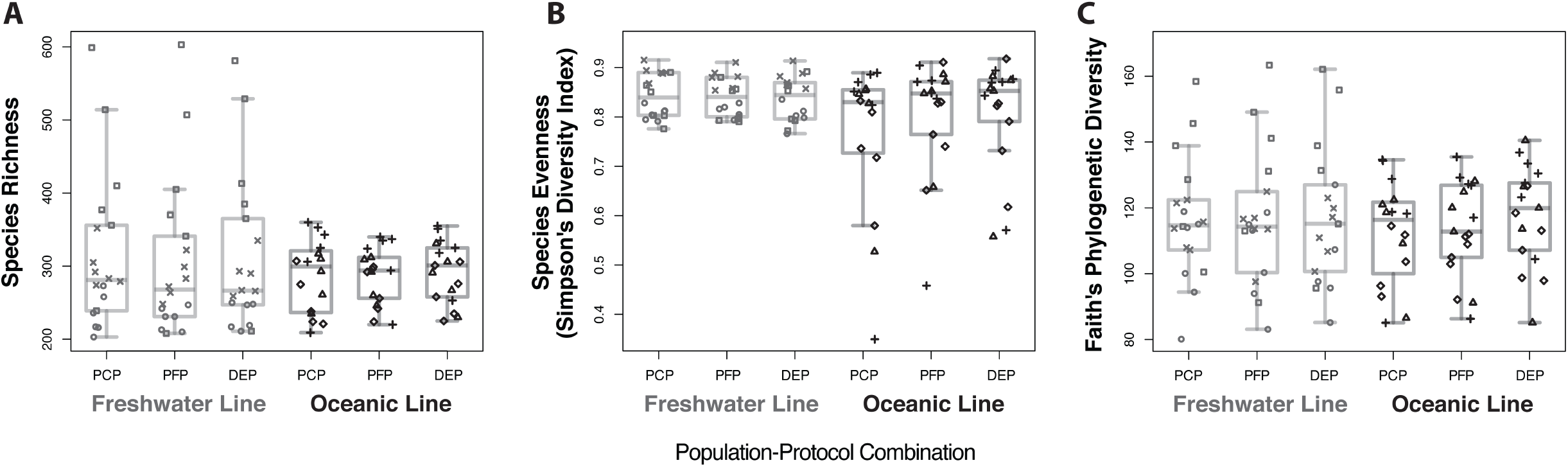
DNA isolation protocol and host population do not strongly influence species richness (**A**), species evenness (**B**), or Faith’s Phylogenetic Diversity (**C**). Boxplots expressing class and evenness distributions for the six population-protocol combinations are overplotted with points representing individual libraries, which are coded by stickleback family. Family 1 = square; Family 2 = circle, Family 3 = “X”; Family 4 = triangle; Family 5 = “+”; Family 6 = diamond.

**Supplementary Figure S6.**
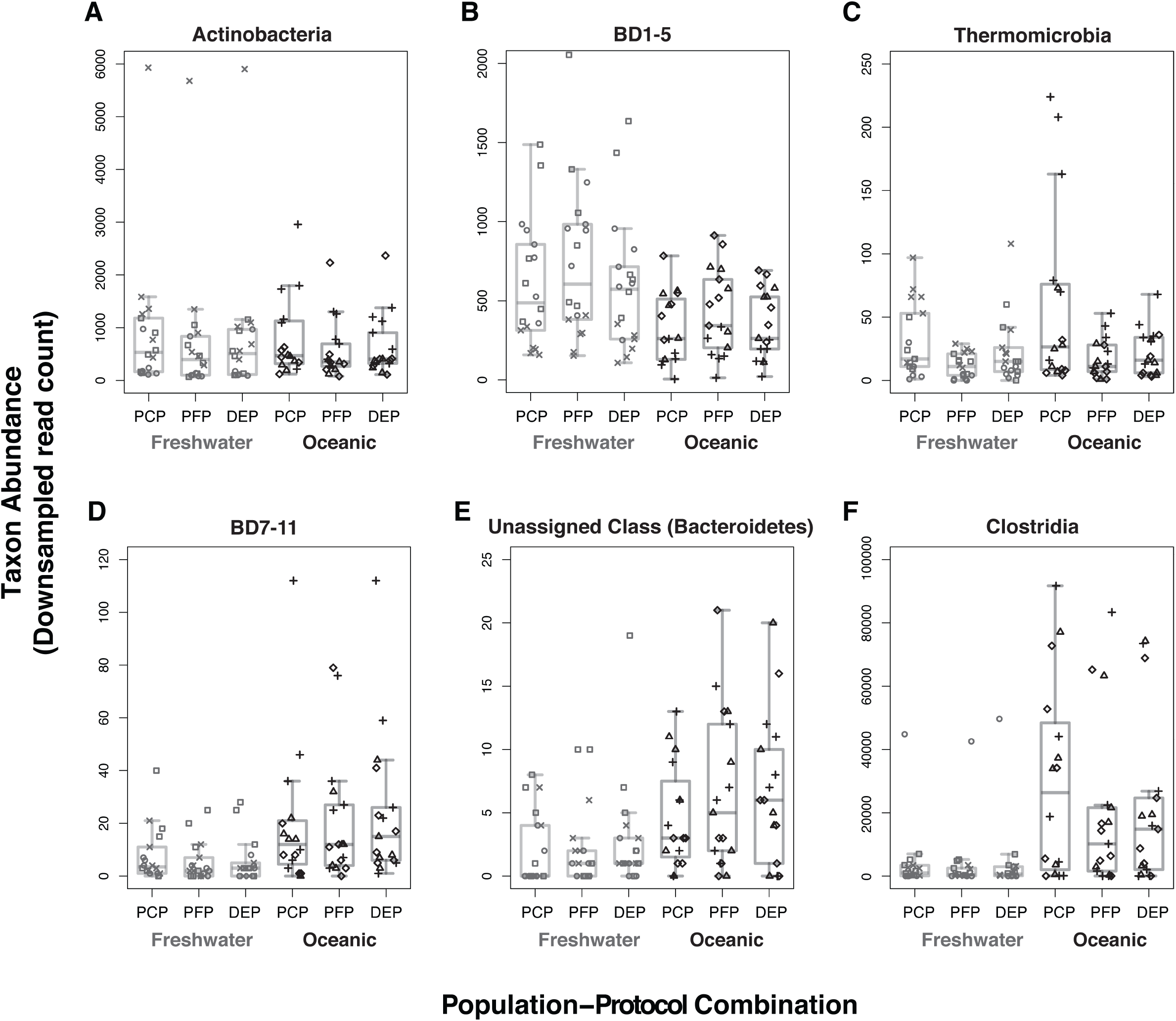
Abundance of individual bacterial classes most likely influenced by DNA isolation protocol (**A-C**) and stickleback population (**D-F**). Boxplots expressing class abundance distributions for the six population-protocol combinations are overplotted with points representing individual libraries, which are coded by stickleback family. Family 1 = square; Family 2 = circle, Family 3 = “X”; Family 4 = triangle; Family 5 = “+”; Family 6 = diamond.

**Supplementary Figure S7.**
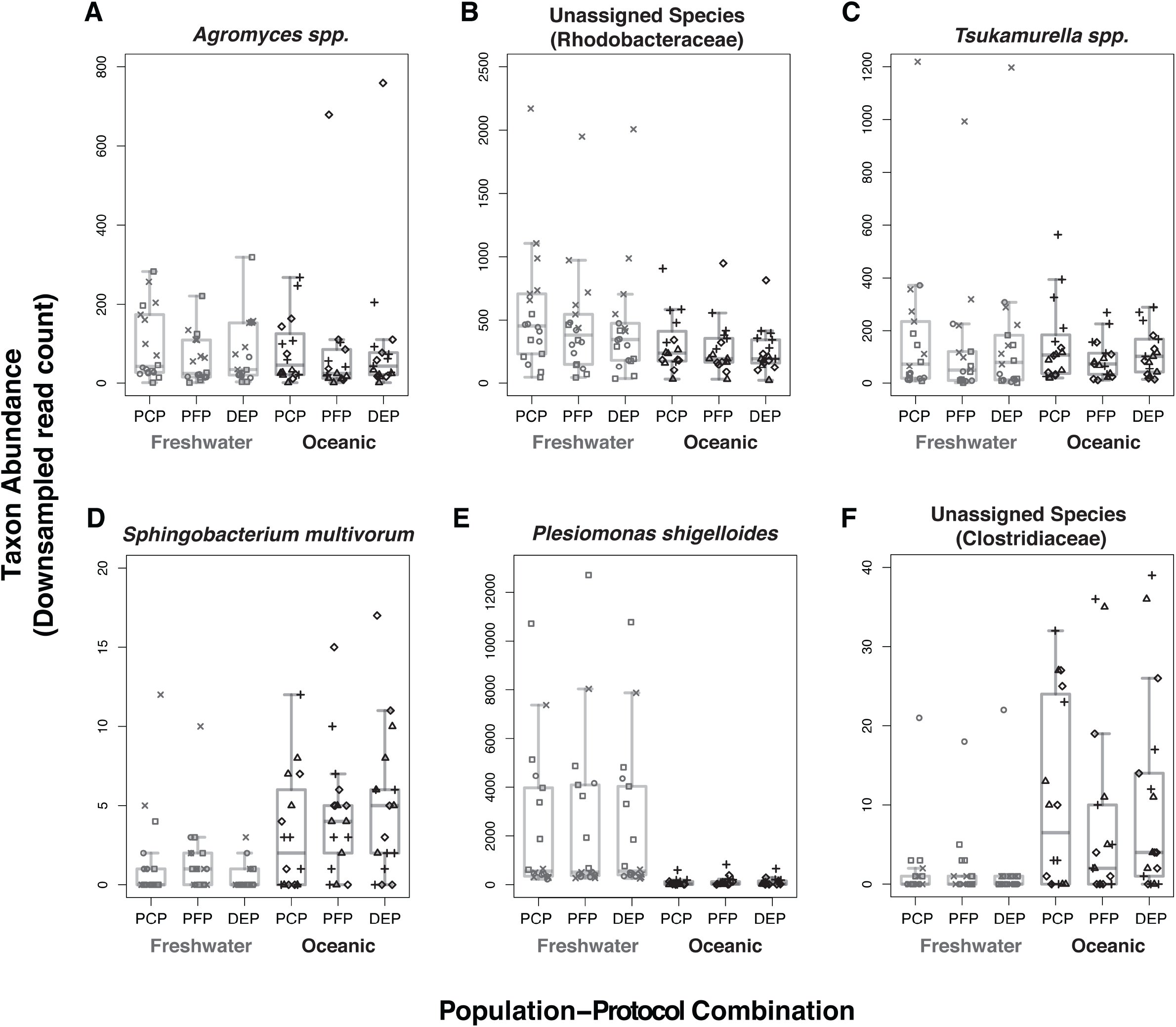
Abundance of individual bacterial species most likely influenced by DNA isolation protocol (**A-C**) and stickleback population (**D-F**). Boxplots expressing species abundance distributions for the six population-protocol combinations are overplotted with points representing individual libraries, which are coded by stickleback family. Family 1 = square; Family 2 = circle, Family 3 = “X”; Family 4 = triangle; Family 5 = “+”; Family 6 = diamond.

**Supplementary Figure S8.**
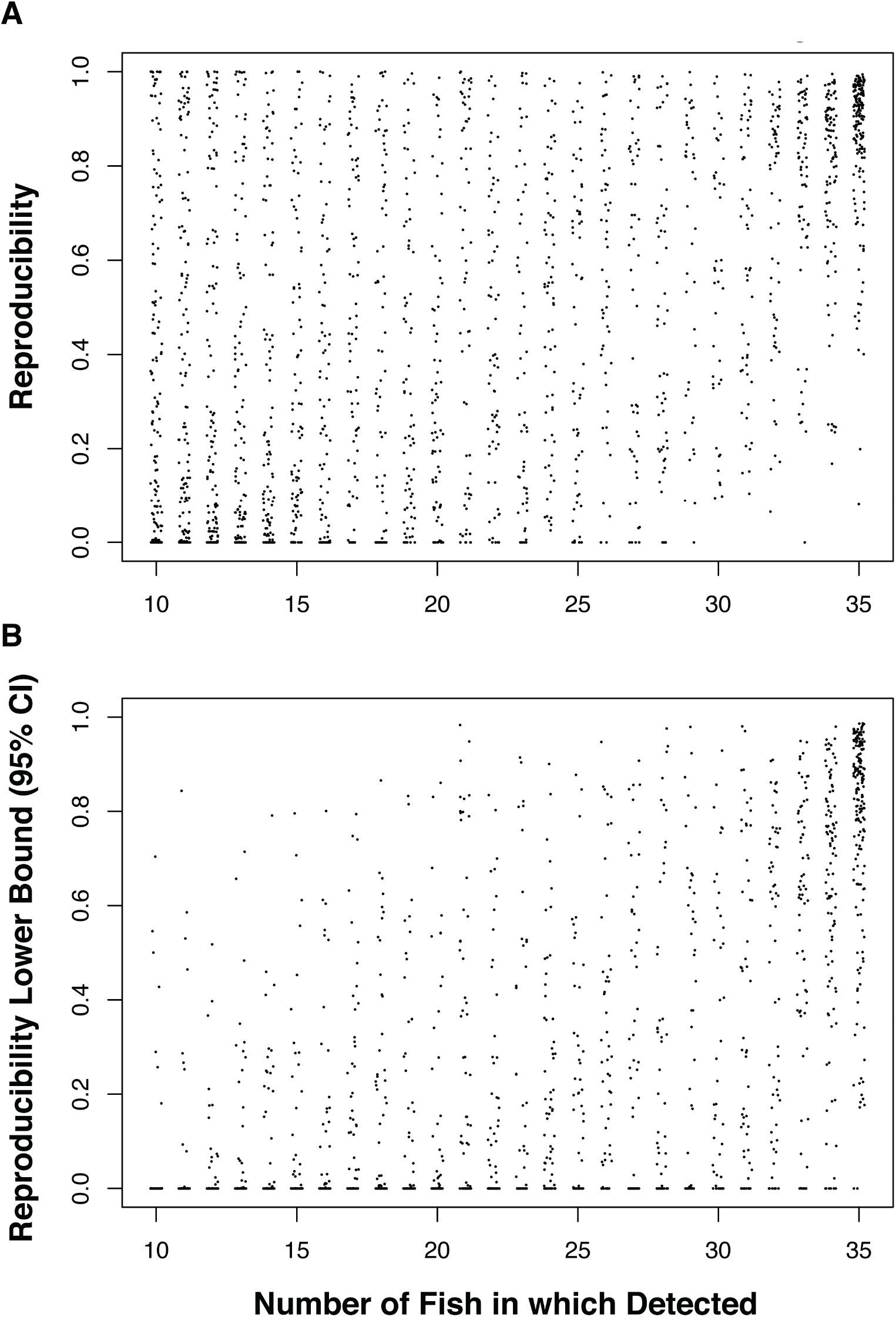
Repeatability of OTU quantification across DNA isolation protocols increases with the number of individuals in which the OTU is detected. **A**. 2278 OTU repeatability estimates plotted against the number of individual stickleback in which present, with x-axis jitter for clarity. **B**. 2278 OTU repeatability lower bound CI estimates plotted against the number of individual stickleback in which present, with x-axis jitter for clarity. A significant, positive shift in repeatability coincided with sampling more than 30 fish in our data set.

**Supplementary Figure S9.**
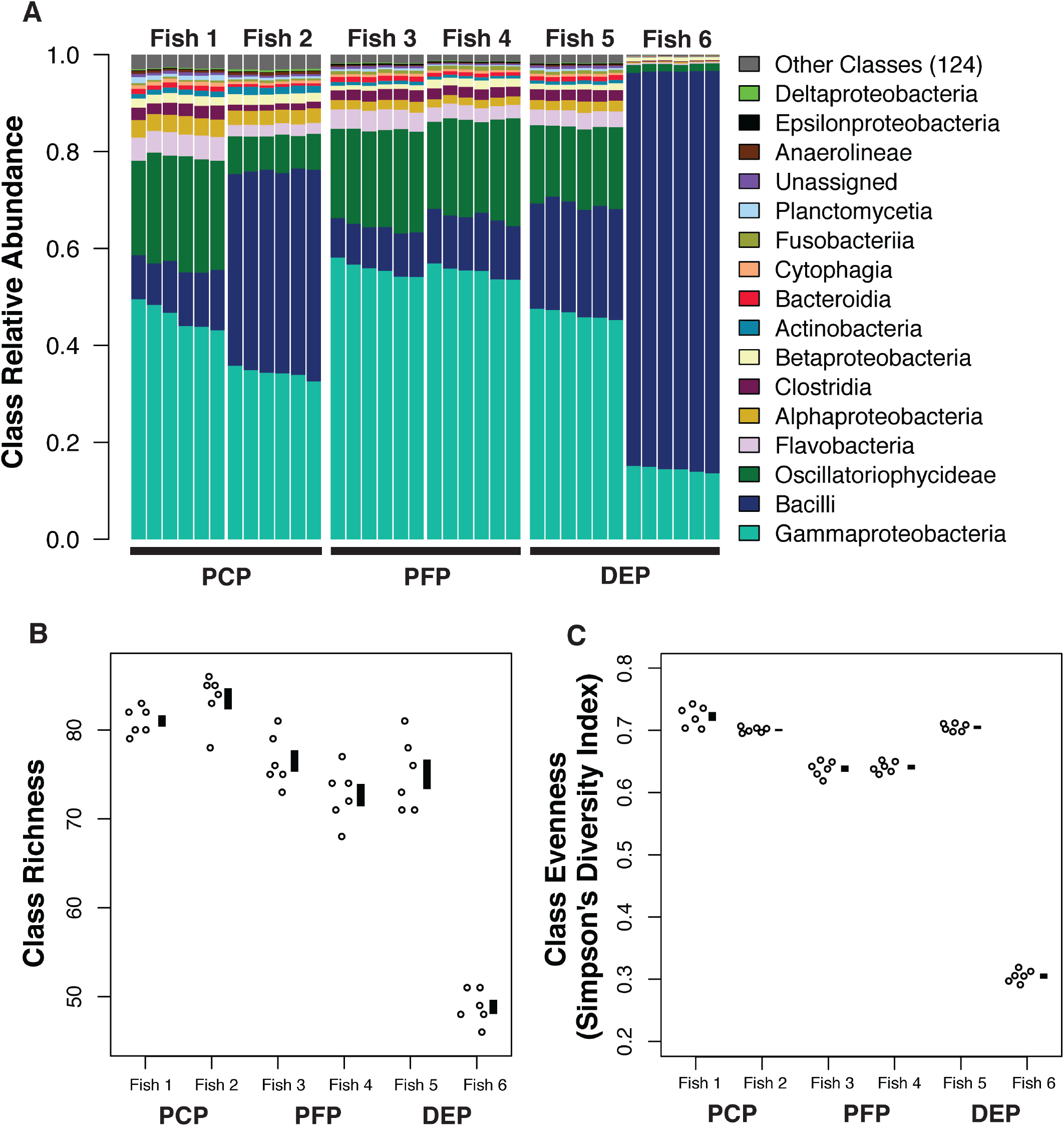
16S sequencing measures class-level microbiota composition and alpha diversity of the stickleback gut with high precision consistently across DNA isolation methods. **A**. Class relative abundances are very similar among technical replicates from the same fish, relative to among-fish differences. Technical replicates with each fish are sorted by mean Gammaproteobacteria abundance. Class richness (**B**) and evenness (**C**) vary substantially among fish, but within-fish (technical) variance is low and similar across the three DNA isolation protocols. Black, vertical bars next to plotted points in **B** and **C** represent mean +/- SEM.

**Supplementary Figure S10.**
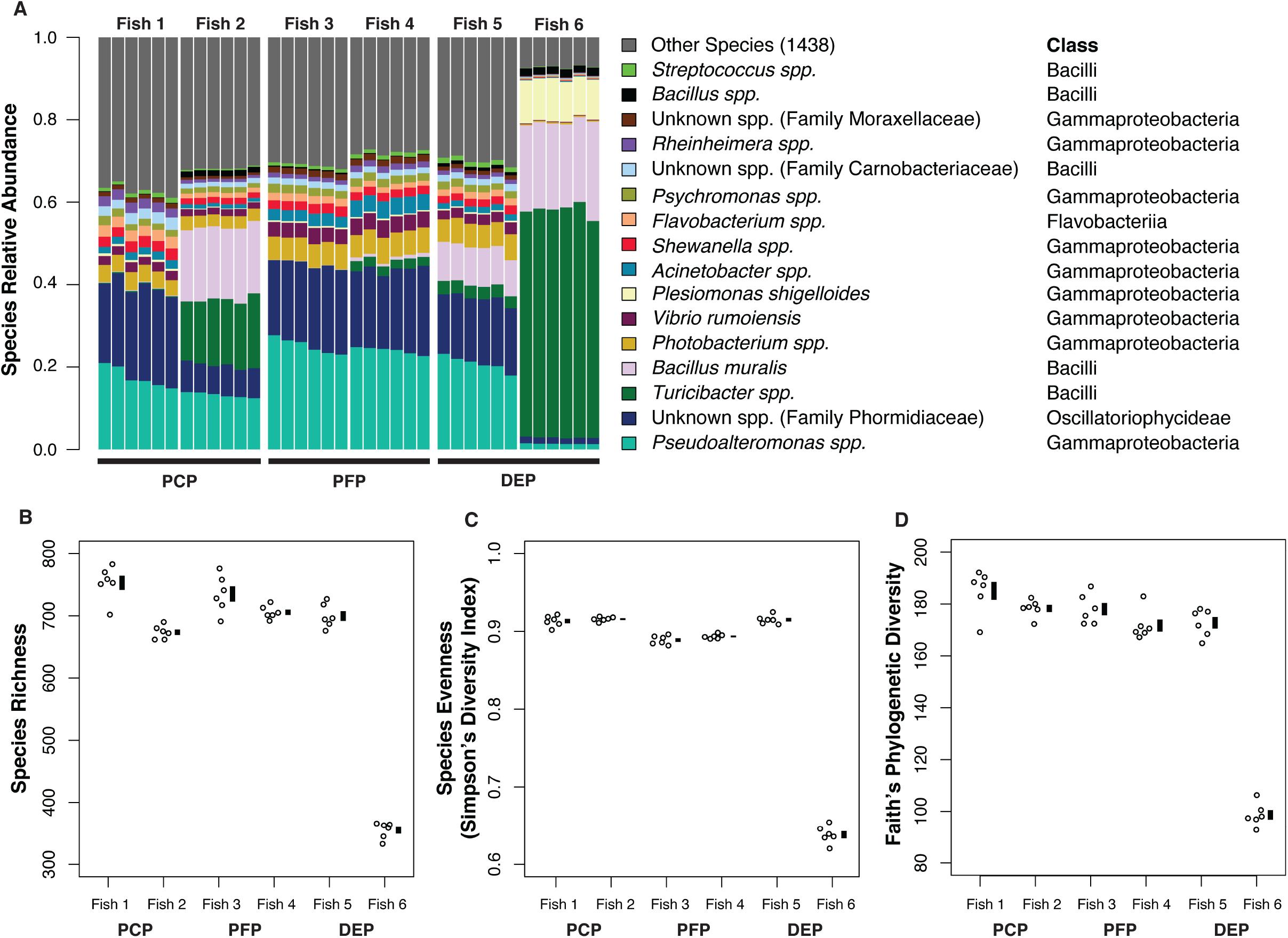
16S sequencing measures species-level microbiota composition and alpha diversity of the stickleback gut with high precision consistently across DNA isolation methods. **A**. Species relative abundances are very similar among technical replicates from the same fish, relative to among-fish differences. Technical replicates with each fish are sorted by mean Gammaproteobacteria abundance. Species richness (**B**) and evenness (**C**) vary substantially among fish, but within-fish (technical) variance is low and similar across the three DNA isolation protocols. Black, vertical bars next to plotted points in **B** and **C** represent mean +/- SEM.

**Supplementary Figure S11.**
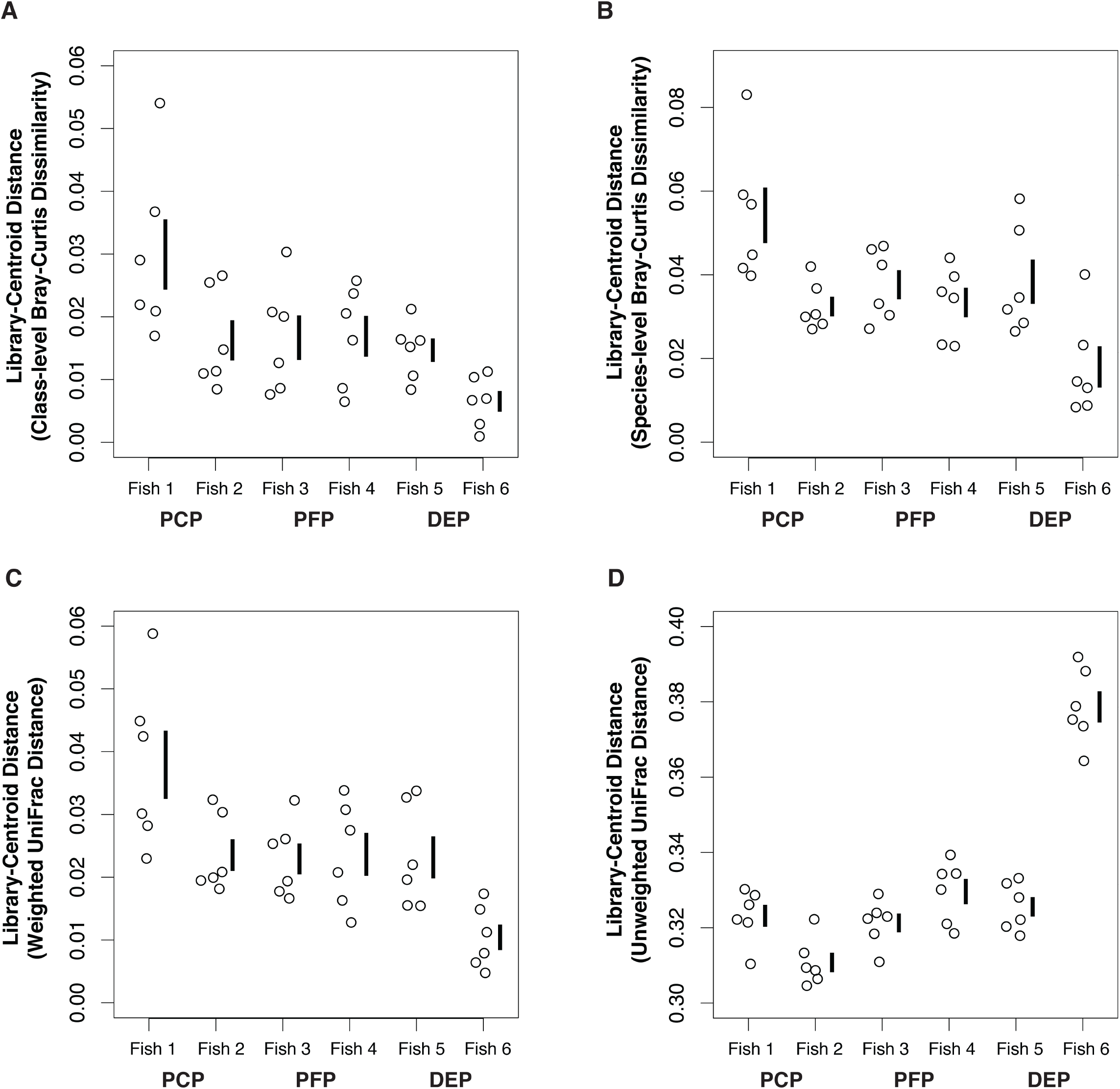
Precision of beta diversity measurements differs among individual hosts, but not consistently among DNA isolation methods. Plotted are multivariate distances between each library and the centroid of its group (fish), which quantify multivariate spread among technical replicates from each fish. Higher y-axis values reflect more spread (lower precision). Black, vertical bars next to plotted points represent mean +/- SEM. Library-centroid distances were calculated based on class- and species-level Bray-Curtis (**A-B**), and weighted and unweighted UniFrac (**C-D**) dissimilarity (see Methods).

### Supplementary Sheet Legends S1A-S1E

<u>Supplementary Sheet S1A</u>: Excel spreadsheet with Permutational Multivariate Analysis of Variance (PERMANOVA) results for protocol, population, family, and family-by-protocol interaction effects on beta diversity, as measured using class- and species-level Bray-Curtis dissimilarity, and weighted and unweighted UniFrac. Shown are results from factorial analyses involving family, protocol, and their interaction, and analysis in which family is nested within population.

<u>Supplementary Sheet S1B</u>: Excel spreadsheet with results from full and reduced lognormal Poisson generalized linear models fit to test effects of Population-by-Protocol Interaction, Population, and Protocol on downsampled counts of individual microbial classes. For each effect tested, reported are the difference in AIC, the difference in BIC, the likelihood ratio test statistic and degrees of freedom, and both uncorrected and FDR-corrected *p*-values. Tests highlighted in pink were associated with a dAIC > 2, a dBIC > 0, and a FDR-corrected *p*-value < 0.1. Population effect tests highlighted in orange were associated with a dAIC > 2, a dBIC > 0, and an uncorrected *p*-value < 0.05.

<u>Supplementary Sheet S1C</u>: Excel spreadsheet with results from full and reduced lognormal Poisson generalized linear models fit to test effects of Population-by-Protocol Interaction, Population, and Protocol on downsampled counts of individual microbial species. For each effect tested, reported are the difference in AIC, the difference in BIC, the likelihood ratio test statistic and degrees of freedom, and both uncorrected and FDR-corrected *p*-values. Tests highlighted in pink were associated with a dAIC > 2, a dBIC > 0, and a FDR-corrected *p*-value < 0.1. Population effect tests highlighted in orange were associated with a dAIC > 2, a dBIC > 0, and an uncorrected *p*-value < 0.05.

<u>Supplementary Sheet S1D</u>: Excel spreadsheet with results (Mantel test *r* statistics and permutation-based *p*-values) from pairwise Mantel tests based on DNA isolation protocol-specific dissimilarity matrices from the reproducibility experiment. Three tests (one for each DNA isolation protocol pair) were performed for each of the four dissimilarity metrics used in this study.

<u>Supplementary Sheet S1E</u>: Excel spreadsheet with results (F-values and *p*-values) from Levene’s Tests comparing variance (univariate) and dispersion (multivariate) among the three different DNA isolation protocols.

